# SLP2/prohibitins aggregates and instability of the PHB complex are key elements in *CHCHD10^S59L^*-related disease

**DOI:** 10.1101/2021.05.31.446377

**Authors:** Emmanuelle C. Genin, Sylvie Bannwarth, Baptiste Ropert, Alessandra Mauri-Crouzet, Françoise Lespinasse, Gaelle Augé, Konstantina Fragaki, Charlotte Cochaud, Sandra Lacas-Gervais, Véronique Paquis-Flucklinger

## Abstract

*CHCHD10* is an ALS/FTD gene, also involved in a large clinical spectrum, that encodes a protein whose precise function within mitochondria is unclear. Here we show that CHCHD10 interacts with the Stomatin-Like Protein 2 (SLP2) to control the stability of the Prohibitin (PHB) complex in the inner mitochondrial membrane. In affected tissues, SLP2 forms aggregates with prohibitins and the instability of the PHB complex results in activation of OMA1 and accelerated OPA1 proteolysis leading to mitochondrial fragmentation, loss of mitochondrial cristae and apoptosis. Abnormal cristae morphogenesis depends on both the PHB complex destabilization leading to MICOS complex instability, *via* disruption of OPA1/Mitofilin interaction, and the activation of PINK1-mediated pathways. We also show that the increase of mitophagy found in both heart and hippocampus of *Chchd10^S59L/+^ mito*-QC mice is PINK1/Parkin-dependent. Thus, SLP2/PHBs aggregates and destabilization of the PHB complex with PINK1 activation are critical in the sequence of events leading to *CHCHD10^S59L^*-related disease.

## Introduction

Mitochondrial diseases, caused by respiratory chain (RC) deficiency, display heterogeneous clinical presentations in terms of age at onset, progression and symptoms. However, pure motor neuron (MN) disorders are rarely seen in these pathologies. Interestingly, we identified a heterozygous mutation (p.Ser59Leu) in the gene encoding Coiled-Coil-Helix-Coiled-Coil-Helix Domain Containing protein 10 (*CHCHD10*) in two families. In the first one, patients developed a mitochondrial myopathy with mtDNA deletions and other symptoms including motor neuron disease (MND), cognitive decline resembling frontotemporal dementia (FTD) and cerebellar ataxia (Bannwarth *et al*, 2014). In the second family, the same p.Ser59Leu variant was responsible for a typical FTD-amyotrophic lateral sclerosis (FTD-ALS) (Bannwarth *et al*, 2014). In both families, the transmission of the disease was autosomal dominant. Mutations in *CHCHD10* were then associated with other clinical presentations, including early-onset mitochondrial myopathies (Ajroud-Driss *et al*, 2015) and cardiomyopathies (Salmon *et al*, 1971; Liu *et al*, 2020). The *CHCHD10* gene was also involved in neurodegenerative disorders such as ALS (Johnson *et al*, 2014; Müller *et al*, 2014), FTD (Sirkis *et al*, 2019), late-onset spinal motor neuropathy (SMAJ) (Penttilä *et al*, 2015) and Charcot-Marie-Tooth disease type 2 (CMT2) (Auranen *et al*, 2015). Among all factors involved in MND, mitochondrial dysfunction has always been recognized as a major player (Cozzolino *et al*, 2015). However, the identification of *CHCHD10* was the first genetic evidence demonstrating that a mitochondrial defect can trigger MND.

The precise function of CHCHD10 in mitochondria is unclear. Fibroblasts of patients carrying the p.Ser59Leu variant (*CHCHD10^S59L/+^*) display RC deficiency, fragmentation of the mitochondrial network, and mitochondrial ultrastructural alterations with loss of cristae (Bannwarth *et al*, 2014). CHCHD10 interacts with Mitofilin/MIC60, a central component of mitochondrial contact site and cristae organizing system (MICOS) complex, the integrity of which is required for the maintenance of mitochondrial cristae (Friedman *et al*, 2015). The expression of the *CHCHD10^S59L^* allele in human cells leads to MICOS complex disassembly and loss of cristae junctions (Genin *et al*, 2016). In mice, *Chchd10* knock-out does not cause disease indicating that *CHCHD10^S59L/+^*-dependent phenotypes are caused by gain-of-function of the mutant protein rather than by loss-of-function (Anderson *et al*, 2019). Recent data in *Drosophila* also support that the *CHCHD10^S59L^* variant is a dominant gain-of-function (i.e., toxic) mutation (Baek *et al*, 2021). In order to understand the biological role of CHCHD10 and to investigate the effects of *CHCHD10* mutations, we generated knock-in (KI) mice with a punctual mutation in the *Chchd10* gene corresponding to the p.Ser59Leu mutation in the human *CHCHD10* gene (Genin *et al*, 2019). *Chchd10^S59L/+^* mice appeared normal at birth but failed to gain weight normally beginning at 10 weeks of age. Around one year of age, they presented the mitochondrial myopathy with mtDNA instability observed in the patients from our original family. Their condition worsened rapidly with severe weight loss and tremors and they developed a fatal mitochondrial cardiomyopathy associated with enhanced mitophagy. At the end-stage of the disease, *Chchd10^S59L/+^* animals displayed neuromuscular junction (NMJ) and MN degeneration with hyperfragmentation of the motor end plate and significant MN loss in lumbar spinal cord. In addition, we observed TDP-43 cytoplasmic aggregates in spinal neurons, corresponding to TDP-43 proteinopathy found in ALS patients (Genin *et al*, 2019). Taken together, these results show that *CHCHD10^S59L/+^* mice are a relevant model for studying CHCHD10 role and pathways dysregulated in *CHCHD10*-associated diseases.

By using both human fibroblasts and mouse models, here we show that SLP2 and prohibitins, major actors in mitochondrial biogenesis, are key elements in the triggering of the pathological cascade leading to *CHCHD10^S59L^*-related disease.

## Results

### CHCHD10 physically interacts with the inner mitochondrial membrane (IMM) scaffold Stomatin-Like Protein 2 (SLP2)

Understanding the role of CHCHD10 requires protein interactome studies. Among the targets identified after immunoprecipitation experiments with CHCHD10-HA protein combined with mass spectrometry analysis, we focused our attention on the IMM protein SLP2. SLP2 is a member of the SPFH (stomatin, prohibitin, flotilin, HflC/K) superfamily, which is composed of scaffold proteins that form ring-like structures and locally specify the protein-lipid composition, and the loss of which impairs mitochondrial functions (Mitsopoulos *et al*, 2015). We confirmed the interaction between CHCHD10 and SLP2 in HeLa cells expressing wild-type or mutant (CHCHD10^S59L-FLAG^) forms. Both were immunoprecipitated by a rabbit polyclonal anti SLP2 antibody (Fig.1A). CHCHD10/SLP2 interaction was further supported by co-immunoprecipitation (co-IP) of the endogenous proteins from whole cell lysates extracted from both control and *CHCHD10*^S59L/+^ human fibroblasts (Fig.1B). Last, we used a proximity ligation assay (PLA Duolink) to analyze *in situ* the proximity between CHCHD10 and SLP2, comparing control and patient fibroblasts. Exclusion of one antibody out of 2 yielded no detectable PLA signal, indicating the specificity of the assay (Sup Fig.1). PLA experiments resulted in a positive labeling in control cells but quantitative analysis revealed a significant decrease of dot number in patient fibroblasts (Fig.1C-D). Western blot analysis showed that expression level of SLP2 was similar in control and patient cells (Fig.1E). Taken together, these results argue for a physical interaction between CHCHD10 and SLP2 that could be disrupted by the expression of the *CHCHD10^S59L^* allele.

**Figure 1.**
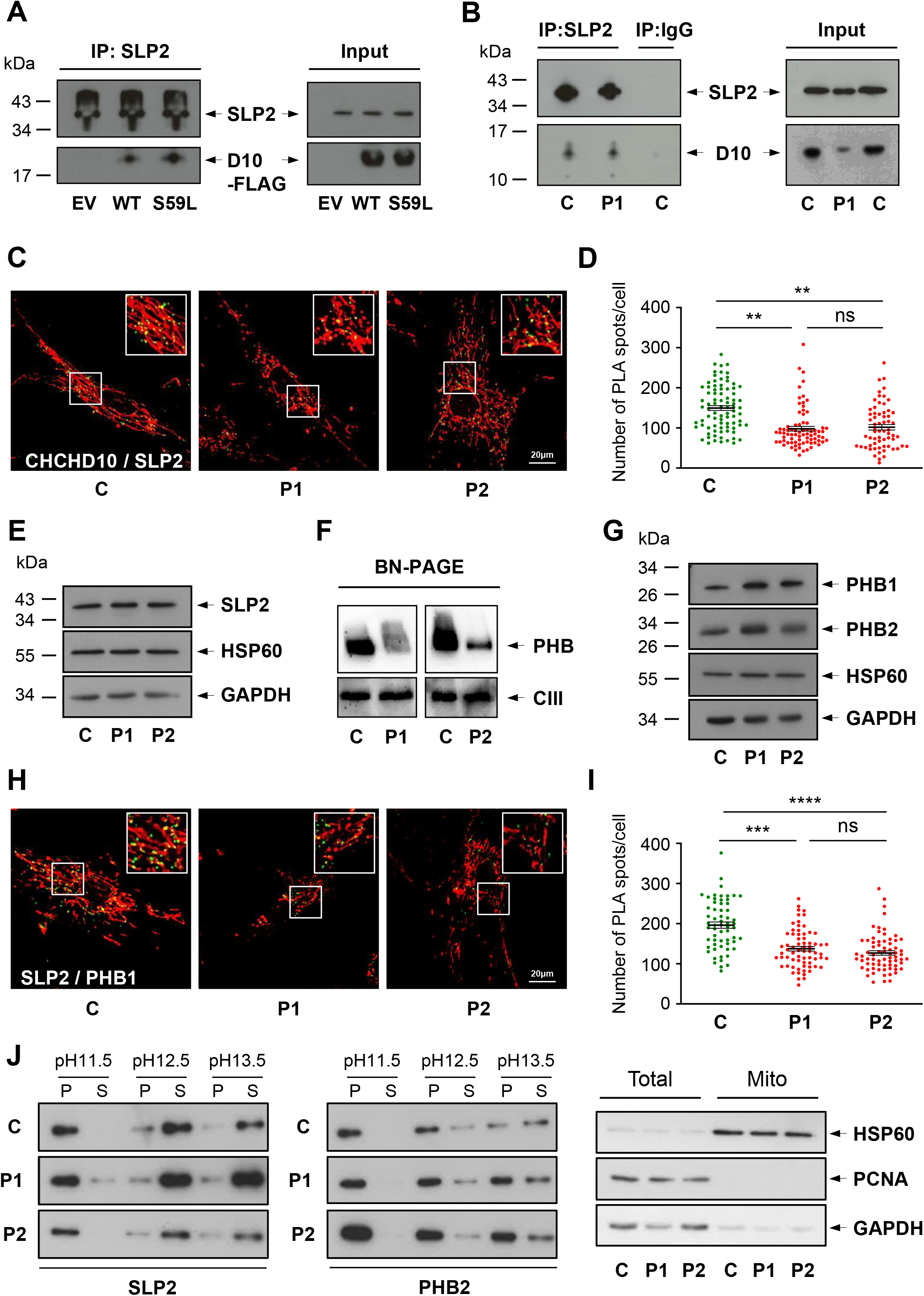
CHCHD10 physically interacts with SLP2 and is involved in the stability of the PHB complex. **A.** Co-immunoprecipitation (IP) experiment in HeLa cells, transfected with an empty vector (EV), wild-type CHCHD10-Flag (WT) or mutant CHCHD10^S59L^-Flag (S59L) constructs, with an antibody against SLP2 revealed with an anti FLAG antibody. **B**. Co-IP of endogenous CHCHD10 and SLP2 in control (C) and patient (P1) fibroblasts. P1 corresponds to patient V-10 harbouring the p.Ser59Leu variant (*CHCHD10^S59L/+^*) (Bannwarth *et al*, 2014). **C**. Duolink proximity ligation assay (PLA) between SLP2 and CHCHD10 in control (C) and patient (P1, P2) fibroblasts observed by confocal microscopy. Mitochondria were stained with MitoTracker. P2 corresponds to patient IV-3 (*CHCHD10^S59L/+^*) (Genin *et al*, 2016). **D.** Duolink spots per cell were quantified for 30 to 40 randomly selected individual cells per each studied fibroblast cell line from two independent experiments. Values are mean ± SEM. **E.** Western blot on control (C) and patient (P1, P2) fibroblast extracts using antibodies against SLP2, HSP60 or GAPDH. **F.** BN-PAGE analysis of mitochondria from control and patient (P1, P2) fibroblasts incubated with an anti PHB2 antibody. Loading control was performed with an antibody against UQCRC2 (complex III, CIII). **G.** Western blot on control (C) and patient (P1, P2) fibroblast extracts using antibodies against PHB1, PHB2, HSP60 or GAPDH. **H.** Duolink PLA between SLP2 and PHB1 in control (C) and patient (P1, P2) fibroblasts observed by confocal microscopy. Mitochondria were stained with MitoTracker. **I.** Duolink spots per cell were quantified for 30 to 35 randomly selected individual cells per each studied fibroblast cell line from two independent experiments. Values are mean ± SEM. **J.** Mitochondrial membranes were extracted from control (C) and patient (P1, P2) fibroblasts at the indicated pHs and separated into pellet (P) and supernatant (S) fractions by centrifugation. Fractions were analyzed by SDS-PAGE and immunoblotting with antibodies against SLP2 and PHB2. To verify the purity of isolated mitochondria, total lysates (Total) and mitochondrial isolates (Mito) were analyzed by immunoblotting using antibodies against HSP60 (mitochondrial protein), PCNA (nuclear protein) or GAPDH (cytosolic protein).

### CHCHD10 is required to maintain the stability of the prohibitin (PHB) complex

SLP2 also interacts and contributes to the stability of prohibitins (PHBs) (Da Cruz *et al*, 2008), which were the first mitochondrial SFPH proteins to be identified (Osman *et al*, 2009). Composed of alternative subunits PHB1 and PHB2, the PHB complex regulates key mitochondrial functions: RC activity, mitochondrial biogenesis, dynamics, apoptosis and mitophagy (for review see Signorile *et al*, 2019). PHBs have been found to be altered in various diseases leading to cellular defects resembling those observed in *CHCHD10*^S59L/+^ cells. We analyzed the PHB complex in both control and *CHCHD10*^S59L/+^ fibroblasts. Blue Native (BN)-PAGE assay revealed a reduction in steady-state PHB levels in patient cells (Fig.1F). Western blot analysis showed that the defect of the PHB complex stability observed in *CHCHD10*^S59L/+^ fibroblasts was not secondary to the proteolysis of PHB1 or/and PHB2 (Fig.1G). To further investigate relationships between PHBs and SLP2 in patient fibroblasts, we used PLA assay to quantitatively assess the proximity between PHB1 and SLP2. As expected, a positive labeling was observed in control cells. However, the significant decrease of dot number in patient fibroblasts was in favor of a disruption of the interaction between SLP2 and prohibitins in *CHCHD10*^S59L/+^ cells (Fig.1H-I). Alkaline extraction of mitochondrial membranes from control fibroblasts at pH 11.5 showed that SLP2 behaves like the integral membrane PHB2, as it is tightly associated with the IMM (Wai *et al*, 2016). At higher pH, SLP2 was mainly recovered in the supernatant fraction due the absence of a transmembrane domain and unlike PHB2 (Fig.1J). We obtained the same results in patient fibroblasts suggesting that SLP2 and PHB2 solubility in the IMM is equivalent in control and patient cells (Fig.1J). Collectively, these data show that CHCHD10 is required to maintain the stability of the PHB complex. They also suggest an alteration of the SLP2/PHB interaction in *CHCHD10*^S59L/+^ cells unrelated to a problem of solubility of these proteins in the IMM.

### SLP2 and PHBs form large aggregates in the hearts of *Chchd10* ^S59L/+^ mice

We analyzed the expression of both SLP2 and prohibitins in the hearts of *Chchd10*^S59L/+^ mice that present with a mitochondrial cardiomyopathy from the age of 3 months. Interestingly, we found a striking accumulation of SLP2 with PHB2 in heart tissue of these mice which was absent in control littermates (Fig.2A). Areas of SLP2/PHB2 accumulation were more visible in the hearts of *Chchd10*^S59L/+^ mice at the end-stage of the disease (Fig.2B). It has been shown that CHCHD10 form large aggregates in *Chchd10*^S59L/+^ hearts (Anderson *et al*, 2019; Genin *et al*, 2019). However, CHCHD10 aggregates do not co-localize with SLP2 foci in *Chchd10*^S59L/+^ heart tissue neither at 3 months nor at the end-stage of the disease (Fig.2C-D). To determine whether SLP2 and prohibitins aggregated in *Chchd10*^S59L/+^ hearts, we used a filter trap assay that captures large protein aggregates. We detected the presence of immunoreactive aggregates for both SLP2 and PHB2 in homogenates from *Chchd10^S59L/+^* mouse hearts but not in hearts from wild type littermates (Fig.2E). CHCHD10 aggregates were also detected in the hearts of the same mice. Therefore, mitochondrial cardiomyopathy presented by *Chchd10^S59L/+^* mice is associated with the accumulation of SLP2/PHBs aggregates which do not co-localize with those involving CHCHD10.

**Figure 2.**
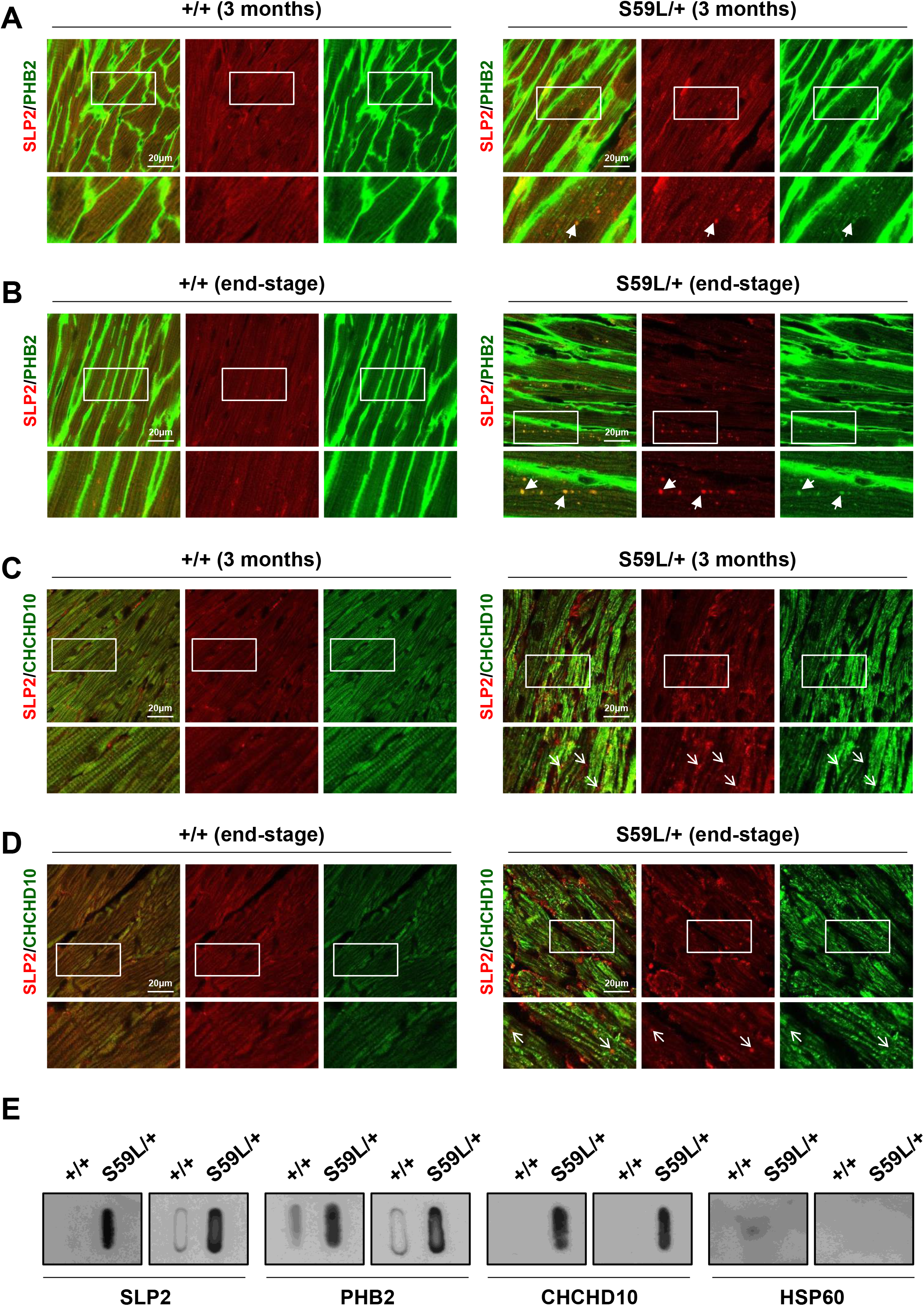
SLP2/PHB2 aggregates in the hearts of *Chchd10* ^S59L/+^ mice. **A-B.** Sections of hearts from WT (+/+) or *Chchd10^S59L/+^* (S59L/+) mice at 3 months of age (A) or at end-stage (B) labeled with antibodies against SLP2 (red) and PHB2 (green). Arrows indicate co-localized puncta. Representative images from 2 independent experiments with 3 *Chchd10^S59L/+^* mice and 3 control littermates. **C-D.** Sections of hearts from WT (+/+) or *Chchd10^S59L/+^* (S59L/+) mice at 3 months of age (C) or at end-stage (D) labeled with antibodies against SLP2 (red) and CHCHD10 (green). Thin arrows indicate non co-localized puncta. Representative images from 2 independent experiments with 3 *Chchd10^S59L/+^* mice and 3 control littermates. **E**. Filter trap analysis of total homogenates from hearts of 2 WT (+/+) or 2 *Chchd10^S59L/+^* (S59L/+) mice at end-stage. Immunoreactive material was detected by immunoblot with SLP2, PHB2 or CHCHD10 antibodies. Anti HSP60 antibody was used as negative control.

### Instability of the PHB complex in the hearts of *Chchd10* ^S59L/+^ mice leads to OMA1-OPA1 cascade activation

We then analyzed the PHB complex *in vivo* and found that its stability was impaired in the hearts of *Chchd10*^S59L/+^ mice at 3 months of age (Fig.3A). Prohibitins and SLP2 control the activity of proteases such as YME1L and OMA1 by keeping them within IMM microdomains. We did not observed an activation of YME1L in the hearts of *Chchd10*^S59L/+^ mice compared to control littermates at 3 months of age (Fig.3B-C). Conversely, OMA1 levels were significantly lower in *Chchd10*^S59L/+^ animals compared to controls (Fig.3D-E). Since OMA1 is degraded shortly after its activation, this result suggests that the instability of the PHB complex triggers OMA1 activation in the hearts of *Chchd10*^S59L/+^ mice (Head *et al*, 2009). The OPA1 (Optic Atrophy type 1) protein is required for mitochondrial fusion, cristae morphogenesis, and apoptosis (Olichon *et al*, 2003; Cipolat *et al*, 2004; Meeusen *et al*, 2006). OMA1 ensures the proteolytic cleavage of OPA1 from long (L-OPA1) to short (S-OPA1) forms at the IMM. Accelerated OPA1 cleavage by OMA1 induces the selective loss of the pro-fusion L-OPA1 forms and the accumulation of the pro-fission S-OPA1 forms leading to mitochondrial fragmentation (Ishihara *et al*, 2006; Baker *et al*, 2014; Quirós *et al*, 2015). In the hearts from *Chchd10^S59L/+^* mice, western blotting analysis revealed a significant decrease of L-OPA1 forms compared to wild type littermates (Fig.3F-G). Independently of its role in mitochondrial fusion, OPA1 is also involved in regulating cristae remodeling and there is a close connection between cristae shape and the assembly of RC complexes (Frezza *et al*, 2006; Cogliati *et al*, 2013). BN-PAGE analysis showed that the stability of several RC complexes (mainly complex IV) was impaired in the heart of *Chchd10^S59L/+^* mice compared to control animals (Fig.3H). Taken together, these results suggest that the instability of the PHB complex found in the hearts of *Chchd10^S59L/+^* mice leads to the activation of the OMA1-OPA1 cascade which is accompanied by a stability defect of the RC complexes.

**Figure 3.**
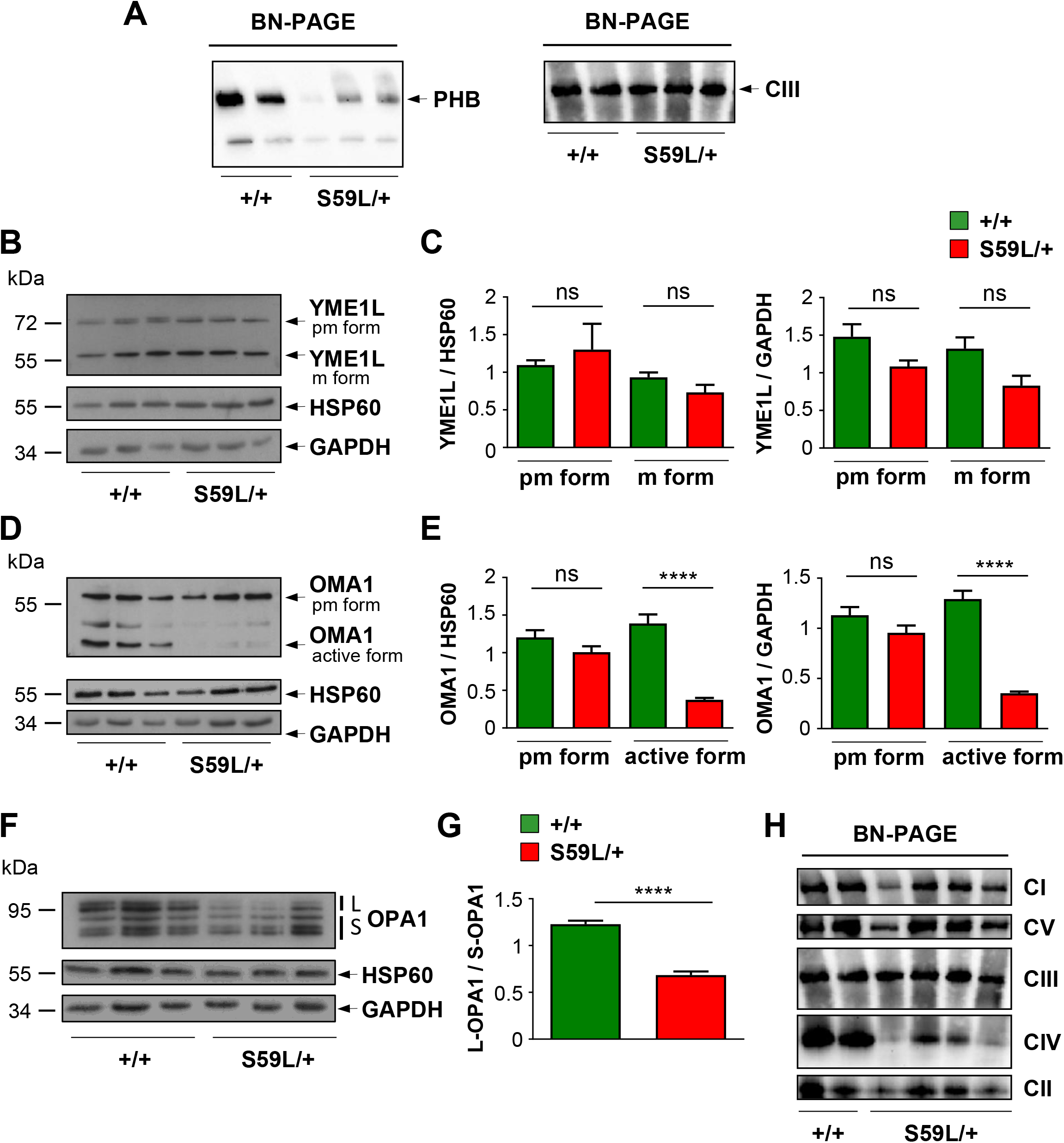
PHB complex disruption, OMA1 activation and increased OPA1 processing in hearts of *Chchd10*^S59L/+^ mice. **A.** BN-PAGE analysis of heart homogenates from *Chchd10^S59L/+^* mice (S59L/+) and control littermates (+/+) performed with an anti PHB2 antibody. Loading control was performed with an antibody against UQCRC2 (Complex III, CIII). **B.** Expression level of YME1L protein analyzed by western blotting in heart tissue from *Chchd10^S59L/+^* mice (S59L/+) and control littermates (+/+). The premature (pm, 75kDa) and mature (m, 63kDa) forms of YME1L are indicated. Antibodies against HSP60 and GAPDH were used for loading control. **C.** Quantification performed from 3 independent experiments with 3 *Chchd10^S59L/+^* mice and 3 control littermates. Values are mean ± SEM. **D.** Representative western blot of OMA1 protein performed with heart lysates from *Chchd10^S59L/+^* mice (S59L/+) and control littermates (+/+). The premature (pm, 60kDa) and active (47kDa) forms of OMA1 are indicated. Antibodies against HSP60 and GAPDH were used for loading control. **E.** Quantification performed from 3 independent experiments with 3 *Chchd10^S59L/+^* mice and 3 control littermates. Values are mean ± SEM. **F.** Expression level of L-OPA1 forms (L : the 2 highest bands) and S-OPA1 (S : the 3 lowest bands) analyzed by western blotting in heart tissues from *Chchd10^S59L/+^* mice (S59L/+) and control littermates (+/+). **G.** Quantification (from 4 independent experiments with 3 *Chchd10^S59L/+^* mice and 3 control littermates) of L-OPA1/S-OPA1 ratios. Values are mean ± SEM. **H.** BN-PAGE analysis of heart homogenates from *Chchd10^S59L/+^* mice (S59L/+) and control littermates (+/+) performed with antibodies against GRIM19 (complex I, CI), SDHA (complex II, CII), UQCRC2 (complex III, CIII), MTCO2 (complex IV, CIV) and ATBP (complex V, CV).

### Depletion of SLP2 or OMA1 compensates fragmentation of the mitochondrial network and abnormal cristae morphology in patient fibroblasts

To investigate whether SLP2 is involved in the sequence of events leading to the fragmentation of the mitochondrial network found in patient cells, we decided to blunt SLP2 expression in *CHCHD10*^S59L/+^ fibroblasts. Transfection with SLP2 siRNA led to SLP2 depletion both in control and patient cells (Sup Fig.2A). We observed that SLP2 depletion rescues the mitochondrial fragmentation found in patient fibroblasts (Fig.4A-B). Moreover, electron microscopy (EM) analysis revealed that mitochondrial morphology and cristae shape were improved after transfection with SLP2 siRNA (Fig.4C-D).

**Figure 4.**
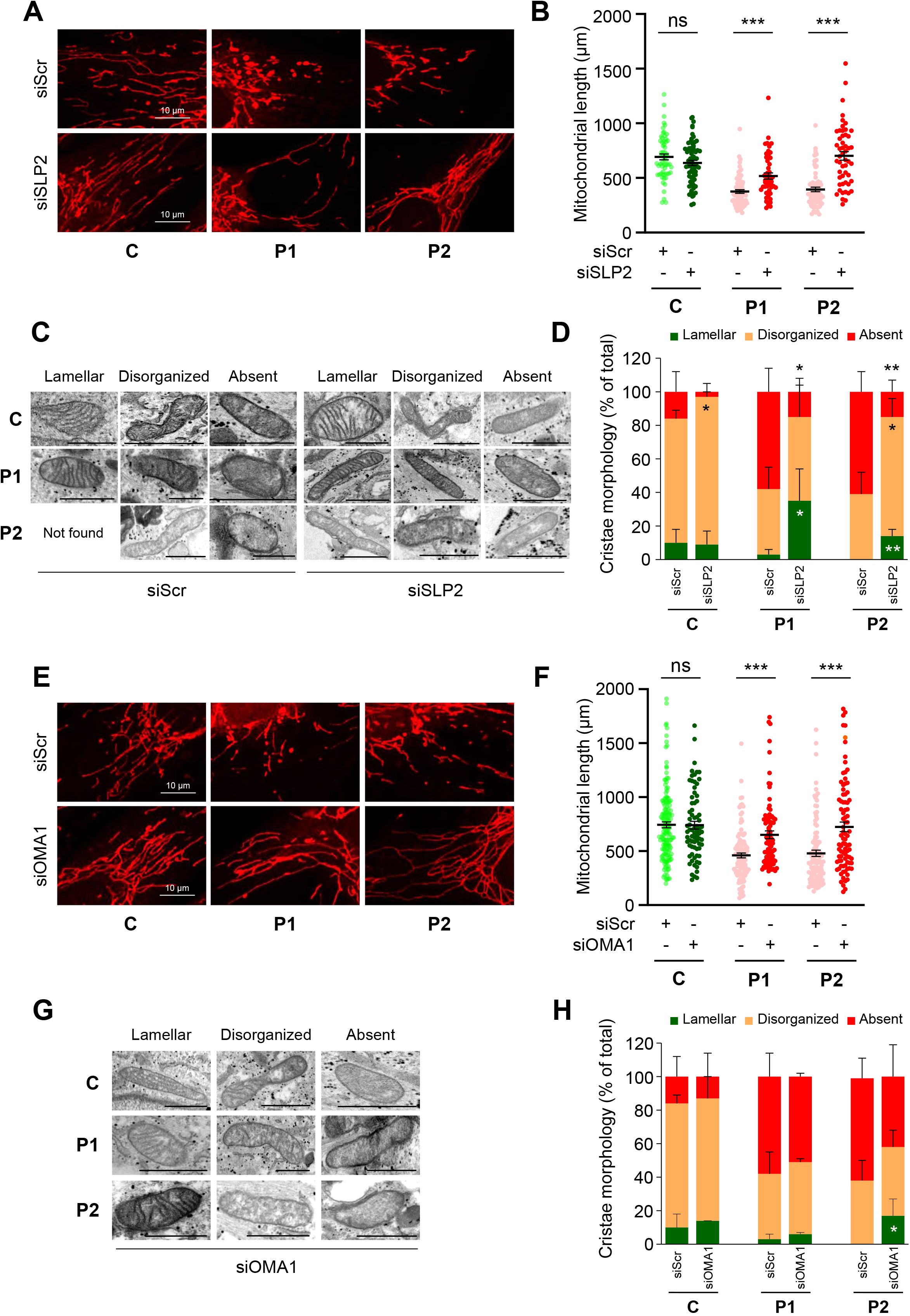
SLP2 or OMA1 depletion rescues mitochondrial network fragmentation and abnormal cristae morphology in *CHCHD10^S59L/+^* fibroblasts. **A.** Control (C) or patient (P1, P2) fibroblasts were transfected with scrambled siRNA (siScr), as negative control, or SLP2 siRNA (siSLP2). Representative images of mitochondrial network from transfected fibroblasts. Mitochondria were stained with MitoTracker. **B.** Quantification of mitochondrial length in transfected fibroblasts. Randomly selected individual 30 to 40 cells per each transfection were analyzed from two independent experiments. Values are mean ± SEM. **C.** Representative images of mitochondrial morphology by electron microscopy (EM), 72h after transfection. Lamellar, disorganized or absent refer to mitochondrial cristae state. Not found corresponds to the absence of mitochondria with lamellar cristae in P2 cells after transfection with siScr. **D.** Quantification of mitochondrial morphology observed in C. n=2 independent experiments (49 to 161 mitochondria per experiment). Values are mean ± SD. **E.** Control (C) or patient (P1, P2) fibroblasts were transfected with scrambled siRNA (siScr), as negative control, or OMA1 siRNA (siOMA1). Representative images of mitochondrial network from transfected fibroblasts. Mitochondria were stained with MitoTracker. **F.** Quantification of mitochondrial length in transfected fibroblasts. Randomly selected individual 30 to 40 cells per each transfection were analyzed from two independent experiments. Values are mean ± SEM. **G.** Representative images of mitochondrial morphology by EM, 72h after transfection with siOMA1. Lamellar, disorganized or absent refer to mitochondrial cristae state. **H.** Quantification of mitochondrial morphology observed in G. n= 3 independent experiments (44 to 132 mitochondria per experiment). Values are mean ± SD.

We then inactivated OMA1 expression in order to confirm the involvement of this protease in *CHCHD10*^S59L^-related phenotypes (Sup Fig.2B). OMA1 depletion was significantly able to drive an elongation of the mitochondrial network in patient fibroblasts (Fig.4E-F). The effect on the mitochondrial cristae disorganization was milder than the one observed after SLP2 depletion (Fig.4G-H). However, these results confirm the role of both SLP2 and OMA1 in the pathological cascade leading to abnormal mitochondrial dynamics and cristae disorganization found in *CHCHD10*^S59L/+^ cells.

### PHB complex instability leads to disassembly of MICOS complex both in control and *CHCHD10^S59L/+^* cells

CHCHD10 physically interacts with Mitofilin/MIC60, a key component of the MICOS complex (Liu *et al*, 2020; Genin *et al*, 2016; Pfanner *et al*, 2014) that is destabilized in *CHCHD10^S59L/+^* fibroblasts (Genin *et al*, 2018). Mitofilin also interacts with OPA1, and together, they control cristae junction number and stability (Glytsou *et al*, 2016; Barrera *et al*, 2016). We performed co-IP analysis to check for OPA1/Mitofilin interaction in *CHCHD10^S59L/+^* fibroblasts. We found that OPA1 was immunoprecipitated by a rabbit polyclonal anti mitofilin antibody as expected in control cells (Fig.5A). However, the level of interaction was lower in patient fibroblasts. This result was confirmed by PLA analysis aimed at studying the proximity between OPA1 and Mitofilin. PLA experiments resulted in a positive labeling in control fibroblasts, but quantitative analysis revealed a significant decrease of dot number in patients cells (Fig.5B-C). Therefore, these results suggest that the disruption of OPA1/Mitofilin interaction plays a role in the lack of stability of the MICOS complex observed in cells expressing the *CHCHD10^S59L^* allele (Genin *et al*, 2016). Then we decided to analyze the proximity between OPA1 and Mitofilin in patients cells after OMA1 depletion (Sup.Fig 3A). Transfection with OMA1 siRNA led to a significant increase of the number of PLA spots in patient fibroblasts suggesting that OMA1 depletion is able to rescue the interaction between OPA1 and mitofilin (Fig.5D-E). Our results also suggest that PHB complex instability, leading to OMA1-OPA1 cascade activation, takes place upstream of MICOS destabilization in the sequence of events leading to loss of cristae junction maintenance. To determine whether PHB complex instability could trigger MICOS complex destabilization, we decided to blunt PHB2 in human fibroblasts. Transfection with PHB2 siRNA led to PHB2 depletion (Sup.Fig 3B). BN-PAGE analysis revealed that it also leads to MICOS destabilization both in control and patient cells (Fig.5F). Collectively, our results show that MICOS complex instability found in *CHCHD10^S59L/+^* cells is a consequence of the PHB complex destabilization.

**Figure 5.**
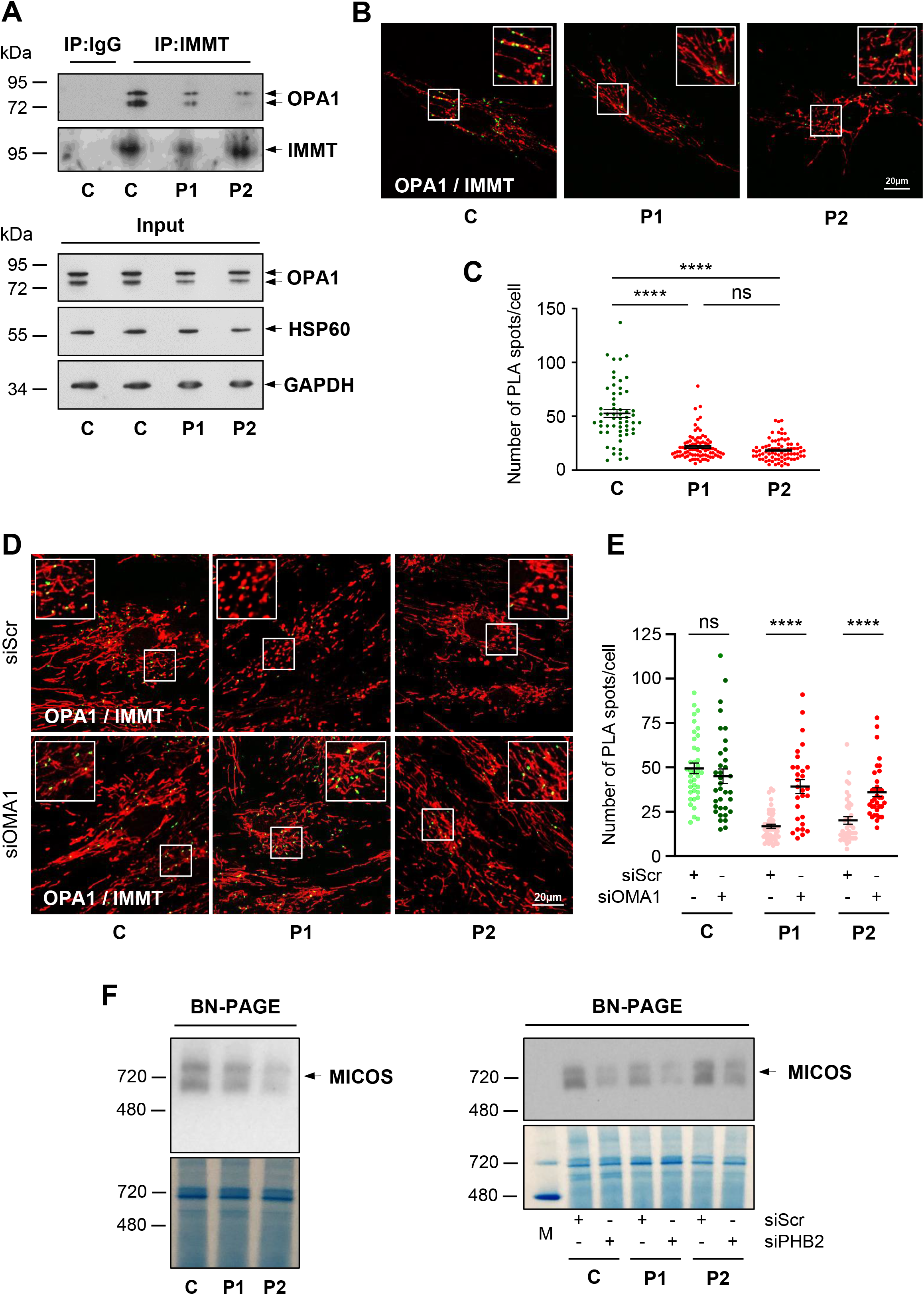
PHB complex stability controls MICOS complex maintenance. **A.** Co-immunoprecipitation (IP) of endogenous mitofilin (IMMT) and OPA1 in control (C) and patient (P1, P2) fibroblasts. **B.** Duolink PLA between IMMT and OPA1 in control (C) and patient (P1, P2) fibroblasts observed by confocal microscopy. Mitochondria were stained with MitoTracker. **C.** Duolink spots per cell were quantified for 30 to 40 randomly selected individual cells per each studied fibroblast cell line from 2 independent experiments. Values are mean ± SEM. **D.** Control (C) or patient (P1, P2) fibroblasts were transfected with scrambled siRNA (siScr), as negative control, or OMA1 siRNA (siOMA1). Duolink PLA between IMMT and OPA1 in control (C) and patient (P1, P2) fibroblasts, transfected with siScr or siOMA1, observed by confocal microscopy. Mitochondria were stained with MitoTracker. **E.** Duolink spots per cell were quantified for 20 to 30 randomly selected individual cells per each studied fibroblast cell line from 2 independent experiments. Values are mean ± SEM. **F.** BN-PAGE analysis showing MICOS state in basal conditions (left panel) or 72h post transfection with scrambled siRNA (siScr), as negative control, or PHB2 siRNA (siPHB2) (right panel) in control (C) and patient (P1, P2) fibroblasts. Mitochondrial proteins from fibroblasts were analyzed by BN-PAGE with an anti mitofilin (IMMT) antibody. Gels were stained with Coomassie brillant blue as loading control. M : NativeMark™ unstained protein standard.

### Abnormal mitochondrial cristae and increased mitophagy found in cells expressing the *CHCHD10^S59L^* allele are PINK1-mediated

The mitochondrial-associated kinase PINK1 plays a critical role in damaged-induced mitophagy (Matsuda *et al*, 2010; Narendra *et al*, 2010). PINK1 is also a crucial factor involved in maintenance of cristae junctions through mitofilin phosphorylation inside mitochondria (Tsai *et al*, 2018). We analyzed the effect of PINK1 depletion on abnormal cristae morphology found in patient cells (Sup. Fig.4A). Electron microscopy analysis revealed that mitochondrial morphology and cristae junctions were improved after transfection with PINK1 siRNA (Fig.6A-B). The degeneration of cardiomyocytes found in *Chchd10^S59L/+^* mice is associated with enhanced mitophagy (Genin *et al*, 2019). To assess the effects of the *Chchd10^S59L^* allele on mitophagy *in vivo*, we used *mito*-QC transgenic mice with a pH-sensitive fluorescent mitochondrial signal, that are widely used to follow mitophagy *in vivo* (McWilliams *et al*, 2016). We crossed *Chchd10^S59L/+^* and *mito*-QC mice to obtain *Chchd10* WT (*Chchd10*^+/+^) and KI (*Chchd10^S59L/+^*) mice on a heterozygous *mito*-QC background. Resultant offspring were born healthy at normal Mendelian frequencies. KI *mito*-QC mice displayed the same clinical evolution than *Chchd10^S59L/+^* animals (Sup. Fig.4B-C). The level of mitophagy, quantified by the number of mitolysosomes, was significantly higher in cardiomyocytes from KI *mito*-QC mice than from WT *mito*-QC animals at 3 months of age. (Fig.6C-D). At end-stage of the disease, mitophagy was also enhanced in KI *mito*-QC cardiomyocytes (Fig.6E-F). The high expression level of the autophagy receptor NDP-52, found in the *Chchd10^S59L/+^* hearts, was consistent with the involvement of the PINK1/Parkin pathway (Fig.6G-H). Immunostaining with Pink1 or Parkin antibodies on heart sections showed a co-localization with mitolysosomes in KI *mito*-QC animals, thus confirming the role of this pathway in the process of mitophagy (Fig.6I). *Phb2* deletion in mouse embryonic fibroblasts (MEFs) renders cells highly susceptible to apoptotic stimuli *via* the selective loss of L-OPA1 isoforms (Merkwirth *et al*, 2008) and pathological conditions of overactive autophagy can lead to cell death (Bloemberg & Quadrilatero, 2019). To assess the effects of the PHB complex instability on cell death *in vivo*, we performed TUNEL staining in *Chchd10^S59L/+^* hearts at end-stage and found evidence of spontaneous apoptosis compared to control tissues (Fig.6J-K).

**Figure 6.**
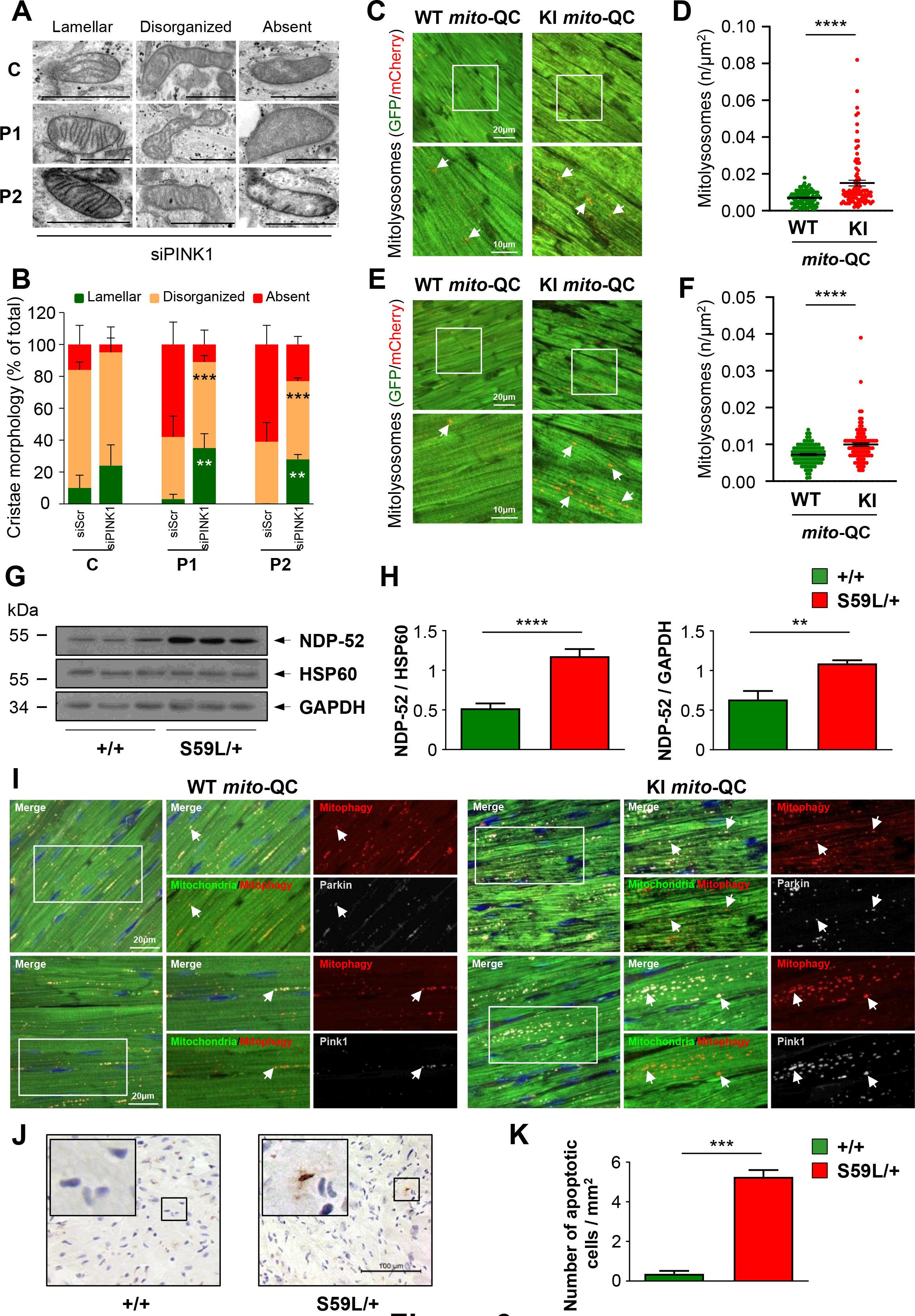
PINK1-mediated abnormal mitochondrial cristae and enhanced mitophagy. **A.** Control (C) or patient (P1, P2) fibroblasts were transfected with scrambled siRNA (siScr), as negative control, or PINK1 siRNA (siPINK1). Representative images of mitochondrial morphology by EM, 72h after transfection with siPINK1. Lamellar, disorganized or absent refer to mitochondrial cristae state. **B.** Quantification of mitochondrial morphology observed in A. n= 3 independent experiments (37 to 137 mitochondria per experiment). Values are mean ± SD. **C, E.** Sections of hearts from WT (*Chchd10^+/+^*) or KI (*Chchd10^S59L/+^*) *mito-*QC mice at 3 months of age (C) or at end-stage (E). Arrows indicate mitolysosomes. **D, F.** Quantification of mitolysosomes observed in C and E, respectively. Data are shown as the mean SEM of 3 independent experiments (3 KI and 3 WT *mito*-QC mice). Thirty fields were randomly selected and analyzed for each animal. **G.** Representative western blot of NDP-52 protein performed with heart lysates from *Chchd10^S59L/+^* mice (S59L/+) and control littermates (+/+). Antibodies against HSP60 and GAPDH were used for loading control. **H.** Quantification (from 3 independent experiments with 3 *Chchd10^S59L/+^* mice and 3 control littermates) of NDP-52/HSP60 and NDP-52/GAPDH ratios. Values are mean ± SEM. **I.** Sections of hearts from WT (*Chchd10^+/+^*) or KI (*Chchd10^S59L/+^*) *mito-*QC mice at end-stage immunolabeled with antibodies against Parkin (upper panels) or Pink1 (lower panels). Arrows indicate co-localization between mitolysosomes and Parkin or Pink1. **J.** TUNEL staining of heart sections from *Chchd10^S59L/+^* mice (S59L/+) and control littermates (+/+). **K.** Quantification of TUNEL positive cells observed in J. Values are mean ± SEM. n= 3 independent experiments with 3 *Chchd10^S59L/+^* mice and 3 control littermates. For each animal, 40 to 68 visual fields were randomly selected to quantify the number of apoptotic cells.

### SLP2/PHBs aggregates with increased mitophagy and apoptosis in the hippocampal areas of *Chchd10^S59L/+^* mice

In order to analyze the effects of the p.Ser59Leu variant in another target tissue of *CHCHD10-*related pathologies, we carried out a BN-PAGE assay from brains of *Chchd10^S59L/+^* mice and control littermates. Since neurological symptoms develop later than heart disorder, we analyzed animals at the end-stage of the disease. Contrary to what had been observed in the hearts of the animals, we did not find any defect in assembly of the PHB complex (Fig.7A). Both YME1L and OMA1 levels were similar in the brains from mutant mice and controls suggesting a lack of activation of these proteases, despite a slight decrease of OMA1 active form (Fig.7B-E). L-OPA1 levels were comparable in the brains from control and mutant animals at the end-stage of the disease (Fig.7F-G). Still contrary to what we observed in the hearts of mice, we did not find any defect in the assembly of the RC complexes by BN-PAGE analysis (Fig.7H). These results suggest either that the neurodegenerative process observed in *Chchd10^S59L/+^* mice is triggered by molecular pathways different from those involved in cardiac damage, or that the activation of the OMA1 cascade is limited to cell types restricted to specific brain areas.

**Figure 7.**
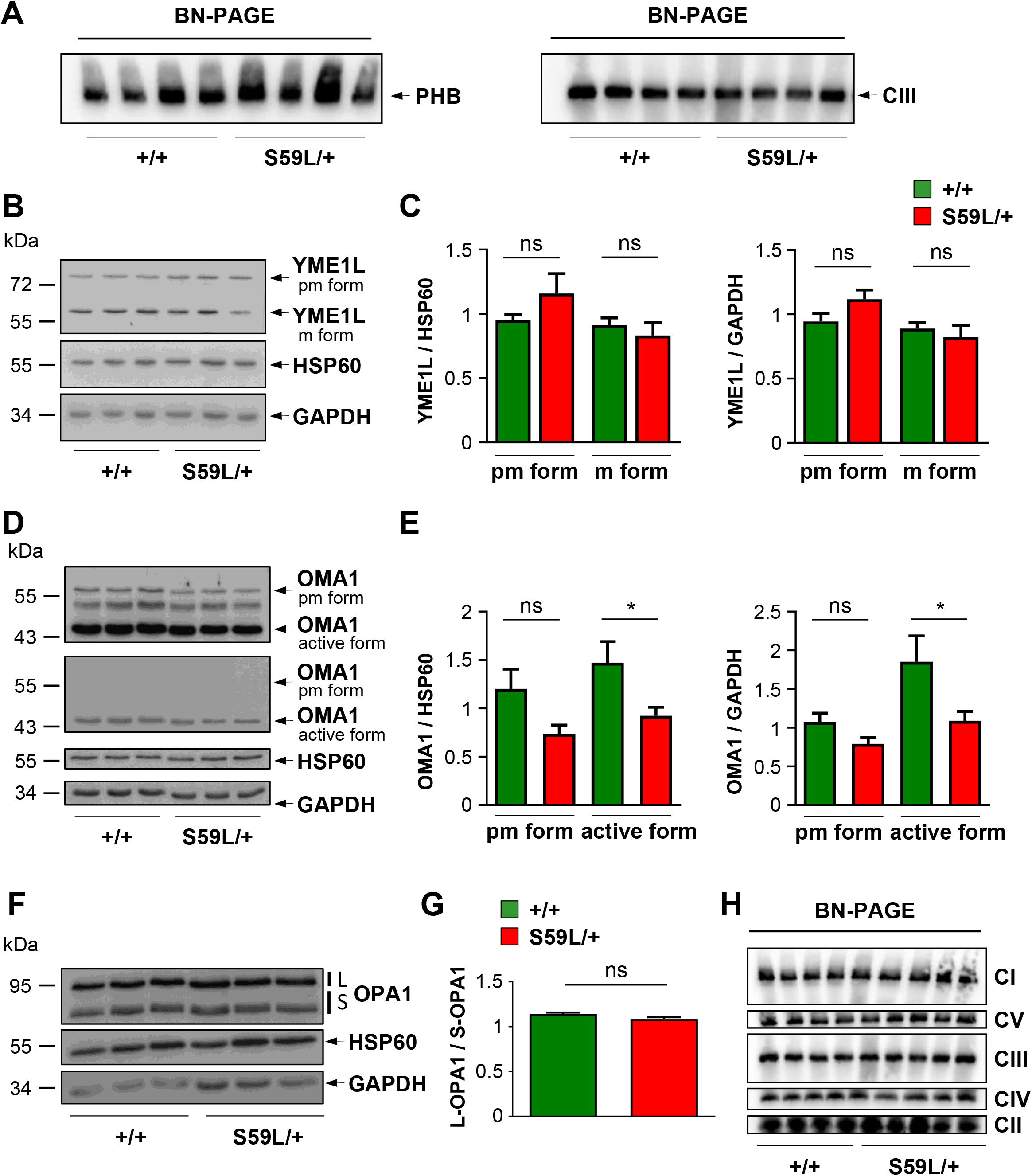
PHB complex integrity without increased OPA1 processing in total brains of *CHCHD10*^S59L/+^ mice. **A.** BN-PAGE analysis of brain homogenates from *Chchd10^S59L/+^* mice (S59L/+) and control littermates (+/+) performed with an anti PHB2 antibody. Loading control was performed with an antibody against UQCRC2 (Complex III, CIII). **B.** Expression level of YME1L protein analyzed by western blotting in brain tissues from *Chchd10^S59L/+^* mice (S59L/+) and control littermates (+/+). The premature (pm, 75kDa) and mature (m, 63kDa) forms of YME1L are indicated. Antibodies against HSP60 and GAPDH were used for loading control. **C.** Quantification performed from 3 independent experiments with 3 *Chchd10^S59L/+^* mice and 3 control littermates. Values are mean ± SEM. **D.** Representative western blot of OMA1 protein performed with brain lysates from *Chchd10^S59L/+^* mice (S59L/+) and control littermates (+/+). The premature (pm, 60kDa) and active (47kDa) forms of OMA1 are indicated. The upper panel corresponds to an overexposure of OMA1 autoradiography. Antibodies against HSP60 and GAPDH were used for loading control. **E.** Quantification performed from 4 independent experiments with 3 *Chchd10^S59L/+^* mice and 3 control littermates. Values are mean ± SEM. **F.** Expression level of L-OPA1 (L) and S-OPA1 (S) forms analyzed by western blotting in brain tissues from *CHCHD10^S59L/+^* mice (S59L/+) and control littermates (+/+). The relative importance of the different isoforms is different from that found in the heart; the highest band being under represented (Akepati *et al*, 2008). **G.** Quantification, from 3 independent experiments experiments (3 *Chchd10^S59L/+^* mice and 3 control littermates), of L-OPA1/S-OPA1 ratios. Values are mean ± SEM. **H.** BN-PAGE analysis of brain homogenates from *Chchd10^S59L/+^* mice (S59L/+) and control littermates (+/+) performed with antibodies against GRIM19 (complex I, CI), SDHA (complex II, CII), UQCRC2 (complex III, CIII), MTCO2 (complex IV, CIV) and ATBP (complex V, CV).

Increased mitophagy is associated with the expression of the *Chchd10^S59L^* allele in the hearts of mice. We took advantage of KI (*Chchd10^S59L/+^) mito-*QC animals to analyse mitophagy in specific brain areas *in vivo*. Interestingly, we observed an increased mitophagy in the hippocampus of KI *mito*-QC mice compared with WT *mito*-QC littermates at the end-stage of the disease (Fig.8A-D). The number of mitolysosomes was significantly higher in dentate gyrus (DG) and in *Cornus Ammonis* CA1 to CA3 (CA) subfields from KI *mito*-QC mice than in the same areas from WT *mito*-QC animals. We also found a significant increase of apoptosis by TUNEL staining in DG and CA from *Chchd10^S59L/+^* mice at the end-stage compared with WT littermates (Fig.8E-H). We then analysed the expression of PHB2 and SLP2 in the hippocampus of *Chchd10^S59L/+^* mice. On the one hand, we found the presence of SLP2/PHB2 aggregates in both the DG and the CA of *Chchd10^S59L/+^* mice which were absent in hippocampal areas of WT littermates (Fig.8I-J). On the other hand, immunostaining with anti PHB2 antibodies revealed totally abnormal labeling both in DG and CA compared to control animals (Fig.8I-J). This pattern was evocative of structural abnormality of PHB complexes. As in patient fibroblasts, we did not find any proteolysis of PHB2 by western blot analysis after microdissection of the hippocampus of mice (Sup. Fig.4D). Therefore, SLP2/PHBs aggregates are associated with increased mitophagy and cell death in the hippocampal areas of *Chchd10^S59L/+^* animals at the end-stage of the disease.

**Figure 8.**
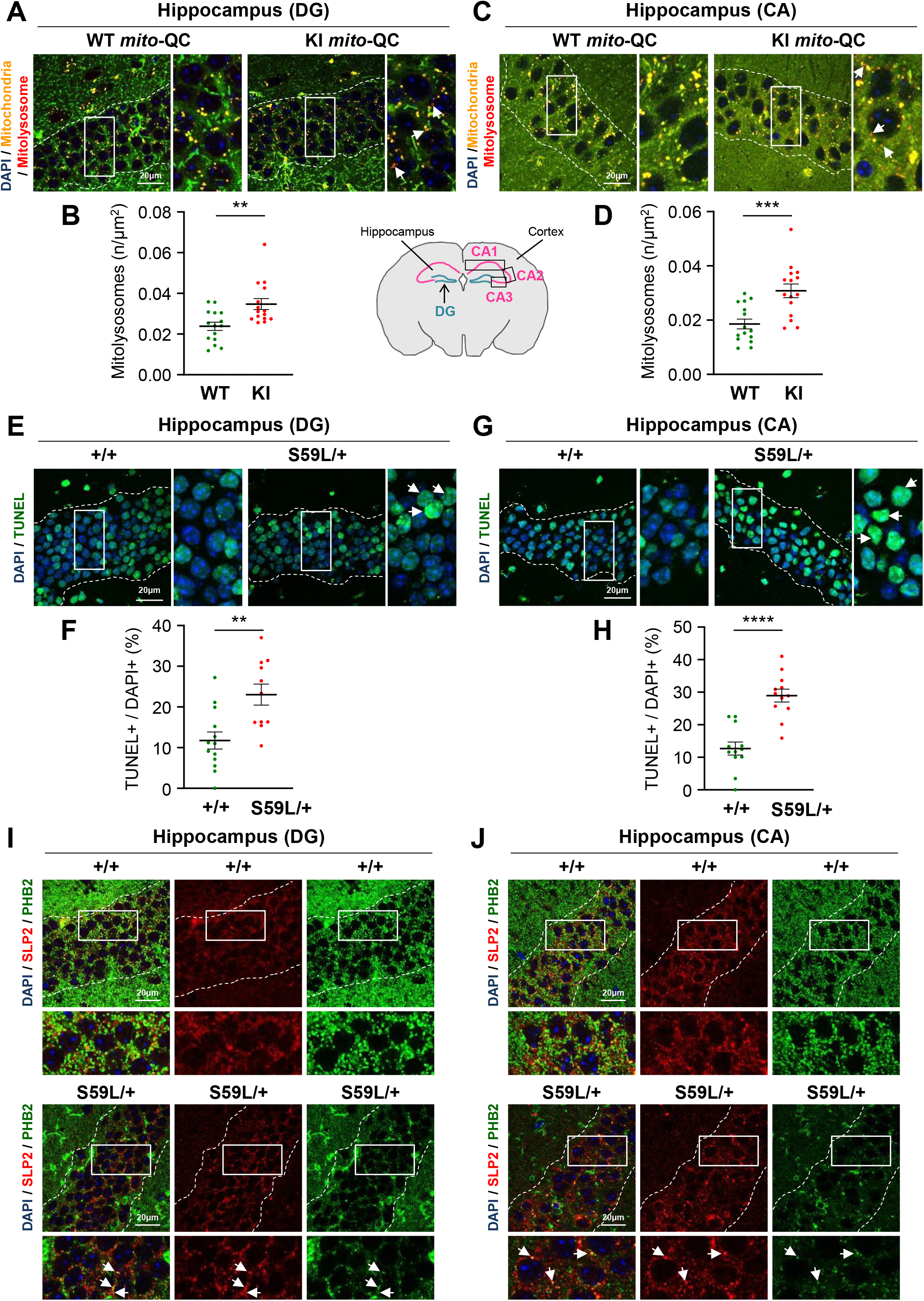
SLP2/PHB2 aggregates with increased mitophagy and apoptosis in the hippocampus of *Chchd10^S59L/+^* mice. **A, C.** Brain sections from WT (*Chchd10^+/+^*) or KI (*Chchd10^S59L/+^*) *mito-*QC mice visualizing hippocampal areas : DG (A) and CA (C) delimited by dotted lines. Arrows indicate mitolysosomes. **B, D.** Quantification of mitolysosomes observed in A and C, respectively. Data are shown as the mean ± SEM of 3 independent experiments (3 KI mice and 3 WT *mito*-QC littermates). Twenty five fields (5 DG or CA per animal and 5 fields per DG or CA1 to CA3) were randomly selected to quantify the number of mitolysosomes. **E, G.** TUNEL staining of brain sections from *Chchd10^S59L/+^* mice (S59L/+) and control littermates (+/+) visualizing DG (E) and CA (G) delimited by dotted lines. Arrows indicate apoptotic (TUNEL+/DAPI+) cells. **F, H.** Quantification of TUNEL positive cells observed in E and G, respectively. n= 3 independent experiments with 3 *Chchd10^S59L/+^* mice and 3 control littermates. For each animal, 4 fields (5 images per fields) from DG or CA1 to CA3 were randomly selected and analyzed for scoring TUNEL-positive cells *versus* total number of nuclei. Values are mean ± SEM. **I-J.** Sections of hippocampal areas : DG (I) and CA (J), delimited by dotted lines, from *Chchd10^S59L/+^* mice (S59L/+) and control littermates (+/+) immunolabeled with antibodies against SLP2 and PHB2. Arrows indicate SLP2/PHB2 co-localized puncta.

## Discussion

Mutations in *CHCHD10* have been found to be responsible for a large clinical spectrum including cardiomyopathy, ALS and/or FTD (Bannwarth *et al*, 2014; Liu *et al*, 2020; Johnson *et al*, 2014; Müller *et al*, 2014). It has been shown that CHCHD10 forms large aggregates with the paralog CHCHD2 protein in the affected tissues and it is very likely that CHCHD10/CHCHD2 aggregation is involved in the pathological process (Anderson *et al*, 2019; Huang *et al*, 2018). Here, we demonstrate that CHCHD10 interacts with SLP2 that has been shown to bind to cardiolipin and to prohibitins (Christie *et al*, 2011). SLP2 forms cardiolipin and prohibitin-enriched microdomains in the IMM which facilitate the assembly of RC complexes (Christie *et al*, 2012). Depleting HeLa cells of SLP2 leads to increased proteolysis of prohibitins and of subunits of the RC complexes I and IV (Da Cruz *et al*, 2008). In affected tissues of *Chchd10^S59L/+^* mice, we found that SLP2 forms large aggregates with prohibitins, which do not co-localize with those involving CHCHD10. Aggregates can lead to the sequestration of the proteins involved which adversely affects their functions in the cell. SLP2/PHBs aggregates are very probably involved in the destabilization of the PHB complex, that we found both in patient fibroblasts and in mouse affected tissues, and that triggers the OMA1-OPA1 activation cascade. It has been shown that the expression of different *CHCHD10* pathogenic alleles both in mouse tissues and in human cells strongly induces the mitochondrial unfolded protein response (mtUPR) leading to an increased expression of mitochondrial proteases and chaperones that control protein folding, assembly and degradation (Anderson *et al*, 2019; Straub *et al*, 2021). CHCHD10/CHCHD2 aggregates may involve sequestration of other proteins essential for mitochondrial homeostasis and trigger the activation of mtUPR (Anderson *et al*, 2019). In *C. elegans*, it has also been reported that the lack of PHB induces a strong mtUPR activation and different observations point to the PHB complex as a key player for mitochondrial quality control and response to mitochondrial stress (reviewed in Hernando-Rodríguez & Artal-Sanz, 2018). Further experiments will be necessary to determine the respective roles of the CHCHD10/CHCHD2 and SLP2/PHBs aggregates in mtUPR triggering and, in the initiation and/or the evolution of the pathological process.

Genetic knockdown of SLP2 or PHB2 results in OMA1 activation (Merkwirth *et al*, 2008; Tondera *et al*, 2009). SLP2 and PHB2 are similar in their function in that they help controlling OMA1 and other AAA-proteases, once more by organizing lipid microdomains in the IMM (Deshwal *et al*, 2020). OMA1 is sequestered in the ring structure formed by the PHB complex. During cell stress, OMA1 is released secondary to PHB complex disruption, converts OPA1 completely into S-OPA1 (pro-fission forms) and triggers mitochondrial fragmentation and cristae disruption (Anand *et al*, 2014; McBride & Soubannier, 2010). Reintroduction of L-OPA1 and not S-OPA1 restores these defects suggesting that S-OPA1 is incompetent for mitochondrial fusion and cristae maintenance (Merkwirth *et al*, 2008). SLP2 and the PHB complex are thus necessary to control the activity of OMA1 and the processing of OPA1. *CHCHD2* variants, involved in Parkinson’s disease (PD), and *CHCHD10* mutations lead to OMA1 activation when overexpressed in human cells (Liu *et al*, 2020). Here, we show that CHCHD10 is required with SLP2 for the maintenance of the PHB complex and that, in our models, OMA1 activation is triggered by the destabilization of this complex. The consequences of OPA1 processing in terms of mitochondrial fragmentation and abnormal cristae morphology are found both in patient fibroblasts and in affected tissues of *Chchd10^S59L/+^* mice. OPA1 physically interacts with Mitofilin/MIC60, a core MICOS component, and together they control the maintenance of mitochondrial cristae (Glytsou *et al*, 2016). More recently, it has been proposed that OMA1 control mitochondrial ultrastructure through its interaction with Mitofilin, independently of OPA1 (Viana *et al*, 2021). We previously found a destabilization of the MICOS complex with abnormal cristae in patient fibroblasts expressing the *CHCHD10^S59L^* variant (Genin *et al*, 2016). Our results place the PHB complex upstream of the MICOS complex in the pathway controlling cristae junction maintenance since PHB2 depletion in control fibroblasts leads to MICOS complex destabilization. In *CHCHD10^S59L/+^* cells, the instability of the PHB complex triggers MICOS complex destabilization through the disruption of OPA1/mitofilin physical interaction. The rescue of mitochondrial endpoints after genetic depletion of SLP2 or of OMA1 in patient fibroblasts confirms the involvement of SLP2 and the OMA1 cascade in the disease.

Our work also highlights the role of PINK1-mediated pathways in our models. In *Drosophila,* rough eye phenotypes induced by overexpression of the human *CHCHD10^S59L^* variant were rescued by *PINK1* depletion (Baek *et al*, 2021). In collaboration with Baek and colleagues, we previously showed that *PINK1* downregulation by siRNA treatment reversed mitochondrial network fragmentation in patient fibroblasts (Baek *et al*, 2021). Here, we show that it also improves morphology of mitochondrial cristae in patient cells, possibly through PINK1/Mitofilin interaction (Tsai *et al*, 2018). In response to stress, aberrant mitochondrial quality control leads to neurodegeneration with induction of the mtUPR triggered by accumulation of misfolded proteins, increased mitochondrial fragmentation and increased mitophagy. These consequences are all found in our models expressing the *CHCHD10^S59L^* variant and it is clear that mitophagy induction is a major phenomenon associated with the expression of the *CHCHD10^S59L^* allele (Baek *et al*, 2021; Genin *et al*, 2019). Upon mitochondrial stress, PINK1 is stabilized on the OMM where it phosphorylates ubiquitin on Ser65 forming pS65-Ub, which recruits and activates Parkin, inducing mitophagy(Pickrell & Youle, 2015). PHB2 also plays a role in the recognition of damaged mitochondria as an IMM receptor required for Parkin-induced mitophagy upon mitochondrial depolarization and rupture of the outer membrane (Wei *et al*, 2017). Thanks to *Chchd10^S59L/+^ mito*-QC mice, we show that the increase of mitophagy in affected tissues is PINK1/Parkin-dependent. Parkin genetic ablation delays disease progression and prolongs survival of SOD1 mouse model of ALS (Palomo *et al*, 2018). Other studies also report that inactivation of PINK1/Parkin-mediated pathways have positive effects in different ALS mouse models, thus suggesting that the activation of PINK1 may be a common mechanism in motor neuron disease (Chen *et al*, 2016; Sun *et al*, 2018). In *CHCHD10^S59L^* - related models, we observed an activation of both PINK1 and OMA1, classically triggered by the same stress, and our results suggest that the link between PINK1 pathway and OMA1 cascade may be PHB-dependent (Yan *et al*, 2020).

Relatively low level mitochondrial stress induces mitophagy to selectively remove damaged mitochondria. However, in response to high-level prolonged stress, mitophagy is followed by cell death (for review see(Bloemberg & Quadrilatero, 2019)). Furthermore, the loss of L-OPA1 forms leads to hypersensitivity to apoptosis (Merkwirth *et al*, 2008). These are probably two reasons why we observed an increased level of apoptosis both in heart and hippocampus of *Chchd10^S59L/+^* mice.

Maintaining a healthy mitochondrial pool is especially important for the viability of long-lived, post mitotic cells such as cardiomyocytes and neurons: two targets in *CHCHD10*-related disease. Cardiac-specific knockout of *Phb2* leads to early-onset severe cardiomyopathy with heart failure (Wu *et al*, 2020). Stress-induced OPA1 processing and mitochondrial fragmentation in mice lead to heart failure which is rescued by OMA1 ablation (Wai *et al*, 2015). Thus, OMA1 has been identified as a potential target to treat cardiomyocyte damage resulting from different etiologies (Acin-Perez *et al*, 2018). The heart phenotype found in *CHCHD10^S59L/+^* mice is fully compatible with the observations resulting from the knockout of *Phb2* and the activation of the OMA1 cascade. It confirms that the inhibition of OMA1 activation may be a therapeutic strategy common to various cardiac pathologies.

It is becoming increasingly evident that mitochondrial alterations are involved in ALS, FTD and other neurodegenerative diseases. These pathologies alter common mitochondrial pathways, the characterization of which is of major therapeutic interest, while affecting distinct neuronal populations. Neuron-specific deletion of *Phb2* in mice induces extensive neurodegeneration (Merkwirth *et al*, 2012). The phenotype is rescued by ablation of OMA1 which delays neuronal loss and prolongs lifespan of the animals (Korwitz *et al*, 2016). A few reports suggest that decreased PHB levels could be involved in the pathogenesis of neurodegenerative diseases by leading to increased apoptosis and neuron loss. A mitochondrial imbalance of PHB1 and PHB2 was reported in intermediate and advanced stages of Alzheimer’s disease (Lachén-Montes *et al*, 2017). A recent work showed that C9orf72, encoded by the most frequently mutated gene in ALS and FTD, is imported in the inter mitochondrial space. C9orf72 recruits the PHB complex to stabilize TIMMDC1, an assembly factor of mitochondrial complex I the function of which is impaired in ALS/FTD patient-derived neurons bearing *C9orf72* mutations (Wang *et al*, 2021). However, to our knowledge, a direct involvement of the instability of the PHB complex has never been reported *in vivo* neither in ALS/FTD nor in other neurodegenerative diseases. We found SLP2/PHBs aggregates in the hippocampus of *Chchd10^S59L/+^* mice at the end-stage of the disease. In addition, immunostaining with anti PHB2 antibodies was in favor of a major destructuring of the PHB complexes in DG and in CA subfields from KI animals compared to WT littermates. This pattern was associated with increased mitophagy and apoptosis in the same hippocampal regions. Different clinical data, especially cognitive and neuromaging findings, suggest that the hippocampus is affected both in ALS and FTD (Abdulla *et al*, 2014; Raaphorst *et al*, 2015; Gami-Patel *et al*, 2021; Gómez-Pinedo *et al*, 2016). The hippocampus is one of the neurogenic niches of the adult brain, which constantly generates neurons throughout life (Eriksson *et al*, 1998). Damage found in *Chchd10^S59L/+^* mice suggests that, in addition to neuron loss, the capacity of cell renewal is impaired. The next steps will be to determine which cell types of the hippocampus are affected and if the molecular pathways identified may be implicated in diseases dependent on other ALS/FTD genes.

In conclusion, we found that CHCHD10 contributes with SLP2 to the maintenance of the PHB complex and indirectly to that of MICOS. In cells expressing the p.Ser59Leu mutation, we show that SLP2/PHBs aggregates and the instability of the PHB complex are key factors which trigger a pathological cascade that leads to cell death in heart and in hippocampus of *Chchd10^S59L/+^* mice (Fig.9). The characterization of the link between the SLP2/PHBs and CHCHD10/CHCHD2 aggregates and the TDP-43 proteinopathy, found in both heart and spinal cord of *Chchd10^S59L/+^* mice, will be essential to determine how the expression of the p.Ser59Leu variant induces aggregation and drives the pathology. *CHCHD10* mutations are relatively rare but the pathways identified by this study could apply to other cardiac and neurodegenerative pathologies and open up common therapeutic possibilities.

**Figure 9.**
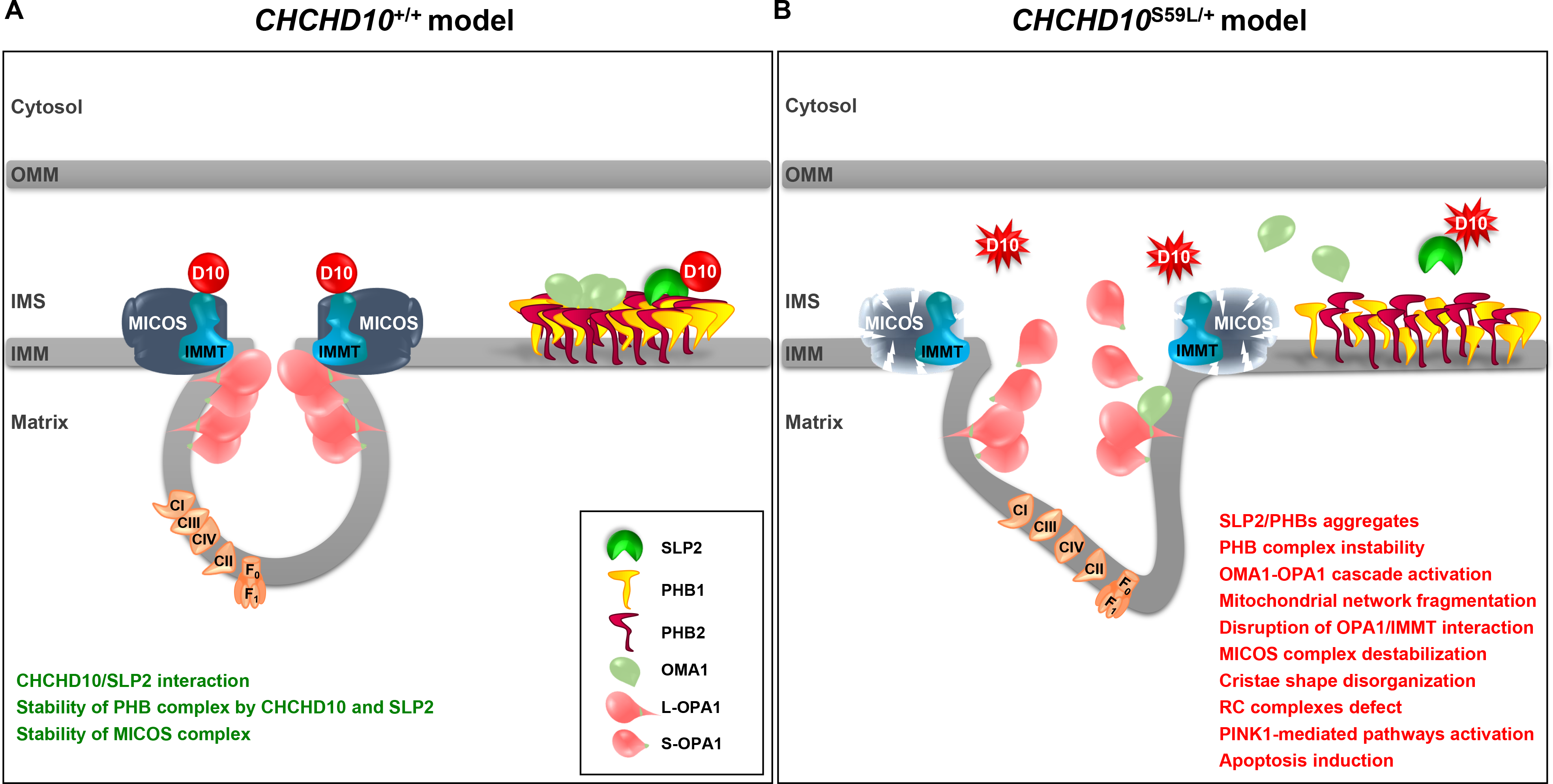
Model for the effects of wild type (A) and mutant (*CHCHD10^S59L^*) (B) alleles in mitochondria. OMM, outer mitochondrial membrane; IMS, intermembrane mitochondrial space; IMM, inner mitochondrial membrane; D10, CHCHD10; IMMT, mitofilin.

## Materials and Methods

### Cell culture

Skin punches were obtained from patients after informed consent. Primary fibroblast cultures were established using standard procedures in RPMI supplemented with 10% Fetal Bovine Serum (FBS), 45 μg/ml uridine and 275 μg/ml sodium pyruvate. Primary fibroblasts and HeLa cells were maintained in DMEM supplemented with penicillin (100U/ml)/streptomycin (0.1mg/ml), 10% FBS. Cell cultures were incubated at 37°C in a humidified atmosphere with 5% CO_2_ in air. For transient transfections, HeLa cells were transfected using Lipofectamine 2000 (Invitrogen) according to the manufacturer’s instructions.

### Co-immunoprecipitation and mass spectrometry experiments

Twenty-four μg of Flag-CHCHD10, Flag-CHCHD10 S59L or Flag alone were transfected in HeLa cells using lipofectamine 2000 (Invitrogen). Twenty-four hours post transfection, cells were subjected to immunoprecipitation assays with 3 mg of proteins and 10μg of mouse anti-FLAG antibody (listed in supplementary table 1). Immunoprecipitated proteins were loaded on a 11% acrylamide-SDS gel. Gel was stained 1 hour with Biosafe Coomassie G250 and destained 45 min in water before proteomic analysis performed at the Proteomics facility of the Institut Jacques Monod, University of Paris Diderot, France.

### Co-immunoprecipitation experiments

Cells were homogenized in solubilization buffer (50 mM Tris-HCl pH7.4, 150 mM NaCl, 0.1% Triton and Halt protease and phosphatase inhibitor cocktail (Thermo Fisher Scientific)) and incubated on ice for 1h. After centrifugation for 5 min at 2600 g and 4°C, cell lysate was incubated with Dynabeads^®^ M-270 Epoxy (Life technologies) for 1 hour at 4°C under gentle rotation. After this pre-clearing step, lysate was incubated 1h at 4°C with gentle rotation with rabbit polyclonal anti-SLP2 antibody, rabbit polyclonal anti IMMT or rabbit IgG (Sigma), previously coupled to Dynabeads^®^ M-270 Epoxy using Dynabeads^®^ Co-Immunoprecipitation Kit (Life Technologies) as described by the manufacturer. After the incubation with the cell lysate, beads were washed three times with solubilization buffer (50 mM Tris-HCl pH7.4, 150 mM NaCl, 0.1% Triton X100 and Halt protease and phosphatase inhibitor cocktail (Thermo Fisher Scientific)). Then, beads were washed with LWB 1× buffer (Dynabeads^®^ Co-Immunoprecipitation Kit (Life Technologies)) and incubated 5 min at RT under gentle rotation. Finally, bound proteins were released with Laemmli buffer. Immunoprecipitated proteins were separated on acrylamide-SDS gels and transferred to PVDF membranes (Millipore). The membranes were probed with rabbit anti SLP2 and mouse anti Flag M2, mouse anti SLP2 and rabbit anti CHCHD10 antibodies or with mouse anti OPA1 and mouse anti IMMT (all listed in supplementary table 1). Then, appropriate secondary antibodies were used at 1/5000 and signals were detected using a chemiluminescence system (Immobilon Western HRP Chemilumiscent substrates, Millipore).

### In situ proximity ligation assay (PLA)

Fibroblasts were seeded in 16-well Lab-Tek chambers slides (Nunc). For mitochondrial staining and after an attachment period of 48h, cells were incubated in a 100nM solution of MitoTracker Red CMXRos (Invitrogen) for 15 min. Then the samples were washed twice with PBS and fixed with PFA 4% for 20 min at 37°C, washed five times with PBS and permeabilised with 2% Triton X-100 for 10 min. After five PBS washes, coverslips were saturated with 5% BSA for 45 min at RT. The following antibodies were used in PLA assay: rabbit anti CHCHD10, mouse anti SLP2, rabbit anti PHB1, rabbit anti OPA1 and mouse anti IMMT (all listed in supplementary table 1). All antibodies were diluted with PBS-BSA 5%. Fibroblasts were incubated in the presence of convenient couple of primary antibodies for 1h at RT. After two PBS washes, incubation with appropriate PLA probe (PLA probe anti rabbit MINUS, PLA probe anti mouse PLUS), hybridization, ligation and amplification were done using the DuoLink *In Situ* Detection Reagents Green (Olink Biosciences) following manufacturer’s instructions. Finally, the samples were mounted on glass slides using Prolong Gold Antifade Reagent (Molecular Probes). Images were captured with a ZEISS LSM 880 confocal laser-scanning microscope and analyzed using Huygens Essential Software™ (Scientific Volume Imaging). Data are represented as mean ± SEM. Negative control experiments (with one antibody omitted) were performed in parallel and checked to result in the absence of PLA signal. Panels obtained with all antibodies used in the manuscript are shown in supplementary figure 1.

### Western blot analysis

The concentration of proteins was determined using the Pierce BCA assay kit (Thermo Fisher Scientific). 2.5-20μg of total protein extracts were separated on acrylamide-SDS gels and transferred to PVDF membranes (Millipore). Specific proteins were detected by using different antibodies listed in supplementary table 1. Signals were detected using a chemiluminescence system (Immobilon Western HRP Chemilumiscent substrates, Millipore). ImageJ was used to quantify protein signals.

### BN-PAGE

BN-PAGE on human fibroblasts were performed using NativePAGE reagent (Invitrogen) according to the standard procedure. Briefly, cells were lysed 15 min on ice in NativePAGE sample buffer 1× containing 1% digitonin or 2% digitonin for the extraction of PHB or MICOS complex, respectively. The lysate was clarified by centrifugation at 20,000xg for 30min at 4°C. Fifteen micrograms of proteins were separated by Blue Native-PAGE Novex 4–16% Bis-Tris gel (Thermo Fisher Scientific). Gels were stained with Coomassie Brilliant Blue. Proteins from the resulting gels were transferred to PVDF membranes (Millipore), and analyzed by western blotting with relevant antibodies. BN-PAGE on mouse tissues were performed with 15μg of mitochondrial respiratory chain complexes obtained by solubilization in a solution of 1.5M aminocaproic acid (Sigma-Aldrich), 75mM Bis-TRIS (Sigma-Aldrich) and 4% dodecyl- b-D-maltoside (Sigma-Aldrich), were separated by Blue Native-PAGE Novex 4– 16% Bis-Tris gel (Thermo Fisher Scientific). Samples were then electroblotted onto a PVDF membrane before sequential incubation with specific antibodies allowing to verify that samples were equally loaded between mutants and controls.

### Alkali extraction

Mitochondria were isolated from fibroblasts using Q-Proteome mitochondria isolation kit (Qiagen) as described by the manufacturer. Mitochondrial membranes were extracted as previously described (Bannwarth *et al*, 2012) with sodium carbonate at the indicated pH. Briefly, intact isolated mitochondria (20 μg) were treated with 0.1M Na_2_CO_3_ (pH 11.5 - 12.5 - 13.5) for 30 min on ice, and then centrifuged at 17,000xg for 15 min at 4°C. Supernatants were retained and pellets were washed once with 200μl of homogenization buffer (250mM sucrose, 1mM EDTA, 20mM Hepes-NaOH pH7.4, plus protease inhibitor) and then resuspended in a 50 μl of homogenization buffer.

### Filter trap assay

Total heart homogenates were treated with 0.5% Igepal for 15 min on ice. After centrifugation, 25 μg of supernatant were filtered through a 0.2μm cellulose acetate membrane (Whatmann) using a Bio-dot microfiltration apparatus. Wells were washed 6 times with 0.1% Tween20 in PBS, membrane was removed from the apparatus, rinsed again 3 times, blocked O/N and immunoblotted with specific antibodies (listed in supplementary table 1).

### siRNA transfection

siRNA transfections were performed using Lipofectamine RNAiMAX reagent (Invitrogen) according to the standard procedure. siRNAs (scramble negative control, human *SLP2, OMA1, PHB2* and *PINK1*) were purchased from Dharmacon (Horizon Discovery). Primary fibroblasts were transfected with 55pmol of siRNA to a final concentration of 55nM and harvested 72h post-transfection for RNA extraction, MitoTracker^®^ staining or protein extraction for western blotting and BN-PAGE analysis.

### RNA extraction, cDNA synthesis and qRT-PCR

Total RNAs were extracted from fibroblasts using TRIzol reagent (ThermoFisher Scientific). Prior to reverse transcription, residual genomic DNA was removed with DNase I (ThermoFisher Scientific). cDNA was then reverse-transcribed using transcription first strand cDNA synthesis kit (Roche Applied Science) with 1 μg total RNA and oligo-dT as primer. All PCRs were performed in triplicate. Quantitative RT-PCR was carried out using SYBR Green master mix (Roche Applied Science) on a Light Cycler LC480. Results were normalized to beta-2macroglubulin (B2M) or 28S genes.

### Mitochondrial network analysis

For mitochondrial staining, cells were incubated with MitoTracker red (Invitrogen) as previously described (Genin *et al*, 2016). The images were deconvolved with Huygens Essential Software™ (Scientific Volume Imaging) using a theoretically calculated point spread function (PSF) for each of the dyes. All selected images were iteratively deconvolved with maximum iterations scored 40 and a quality threshold at 0.05. The deconvolved images were used for quantitative mitochondrial network analysis with Huygens Essentiel Software™ with the following standardized set of parameters: threshold = 10-15%, seed = 10% and garbage = 10. The quantitative data were further analyzed in Microsoft Excel and GraphPad Prism 5 (GraphPad Software). Mitochondrial network length was quantified for 30 to 40 randomly-selected individual cells. Data are represented as mean ± SEM.

### Electron microscopy

Cells were seeded on 24 wells/plate, fixed with 1.6% glutaraldehyde in 0.1 M cacodylate buffer (pH 7.4) for 2 hours, rinsed and postfixed in 1% osmium tetroxide and 1% potassium ferrocyanide in 0.1M cacodylate buffer before to processed for ultrastructure as previously described (Bannwarth *et al*, 2014). The protocole for contrasting sections was slightly modified, using a 5% Lanthanide salt solution (a mixture of Gadolinium acetate and Samarium acetate) for 7 minutes instead of uranium acetate.

### *Chchd10^S59L/+^* mice

All animal procedures were approved by the Institutional Animal Care and Use Committee at the University of Nice Côte d’Azur (Nice, France) and by the Ministère de l’Education Nationale, de l’Enseignement Supérieur et de la Recherche (MESR agreement: APAFIS#5870-2016061017306888). *CHCHD10*^S59L/+^ mice were generated in a previous work (Genin *et al*, 2019).

### *Mito*-QC mice

All animal procedures were approved by the Institutional Animal Care and Use Committee at the University of Nice Côte d’azur (Nice, France) and by the Ministère de l’Education Nationale, de l’Enseignement Supérieur et de la Recherche (MESR agreement: APAFIS#21221-201906211102139). *Mito*-QC mice (C57BL/6J genetic background) have been crossed with *Chchd10^S59L/+^* mice (C57BL/6N genetic background) to generate *Chchd10^S59L/+^* (KI) or *Chchd10*^+/+^ (WT) *mito*-QC mice used in this study.

“*Mito*-*QC Counter*”, a semi-automatic macro for FIJI (ImageJ software), was used to assess mitophagy, as described in Montava-Garriga *et al*. (Montava-Garriga *et al*, 2020).

### Immunohistofluorescence

Mice were treated with buprenorphine (0.1mg/kg) and deeply anesthetized with xylazine/ketamine (15mg/kg and 70mg/kg respectively). Then, they were transcardially perfused with 0.9% NaCl, followed by 4% paraformaldehyde (PFA). Tissues were post-fixed in 4% PFA for 4 hours and then cryoprotected in 20% sucrose for 24 hours. Samples were snap-frozen in isopentane after cryoprotection with Cryomatrix (Thermo Scientific). For brain, 40-μm-thick free-floating serial coronal sections were sliced using cryostat. For heart, 30-μm-thick free-floating serial sections were sliced using cryostat.

For immunostaining, sections were incubated in blocking buffer (0.3-0.5% Triton X-100, 5% normal goat serum, 3-5% bovine serum albumin (BSA)) for 1 hour at RT. Floating sections were stained with primary antibodies diluted in blocking buffer O/N, at 4°C. The following primary antibodies were used: anti rabbit CHCHD10, anti mouse Parkin, anti mouse PHB2, anti mouse PINK1, anti mouse SLP2 or anti rabbit SLP2. For immunstaining with anti mouse SLP2 antibody, antigen retrieval was performed with Tris pH9.0 (Vector Laboratories) for 15 minutes at 95°C and 15 minutes at RT. The references of antibodies and the concentrations used are listed in Supplementary Table 1. After several washings in PBS/Triton 0.1%, sections were incubated for 1 hour, at RT, in a solution (PBS/Triton 0.3%) with the corresponding secondary antibodies conjugated to Alexa Fluor 488, 547 or 647 (Thermo Fisher Scientific) (listed in Supplementary Table 1). Sections were rinsed in PBS/Triton 0.1% and PBS, and incubated with 4′,6-diamidino-2-phenylindole, dihydrochloride (DAPI) (2μg/ml) (Life Technologies) for 5 minutes at RT. Finally, sections were mounted in Vectashield Hard Set medium (Vector Laboratories). All images were captured with a ZEISS LSM 880 confocal laser-scanning microscope or ZEISS LSM 800 confocal laser-scanning microscope.

### TUNEL assay

Brain tissues from animals perfused with PFA 4% were cut into 40μm sections and stained with the *In Situ* Cell Death Detection Kit, Fluorescein (Roche). Briefly, sections were treated with antigen unmasking solution H3300 (Vector laboratories) for 20 min at 92°C, washed in PBS and blocked for 60 min in Tris-HCL 0.1M pH7.5, 3% BSA, 20% normal goat serum, 0.1% Triton. Sections were mounted onto glass slides and reaction mix containing fluorescein-dUTP and terminal deoxynucleotidyl transferase (TdT) was applied for 60 min at 37°C. A set of sections was incubated in the absence of TdT as a negative control. After PBS washes and counterstain with DAPI 2μg/ml for 5 min, sections were washed again in PBS. Images were captured with a ZEISS LSM 800 confocal laser-scanning microscope.

Heart tissues were collected and frozen in liquid nitrogen. TUNEL assay was carried out on cryosections (15μm thickness) using Apoptag Peroxidase *In Situ* Apoptosis Detection Kit (Millipore) as instructed by the manufacturer. Apoptotic cells were labeled brown under DAB staining (Vector laboratories) and sections were lightly counterstained with Mayer Hematoxylin and observed with microscope Leica DM4000B.

### Statistical Analysis

Statistical analysis was done with Student’s *t*-test, Mann-Whitney’s test or Log-rank (Mantel-Cox) test. The quantitative data were analyzed in Microsoft Excel and GraphPad Prism (GraphPad Software). Data are means ± SD or ± SEM. “n” represents number of mice or cells per experiment. The data from *Chchd10*^S59L/+^ mice or KI *mito*-QC mice were compared to the controls littermates. P-value = *<0.05, **<0.01, ***<0.001 and ****<0.0001.

## Acknowledgements

We acknowledge Ian Ganley for the gift of the *mito*-QC mice, and Thibault Léger and the Proteomics facility of the Institut Jacques Monod, University of Paris Diderot for mass spectrometry analysis. We also thank Christelle Boscagli for technical help.

We gratefully acknowledge the IRCAN’s Molecular and Cellular Core Imaging (PICMI) facility, supported financially by FEDER, Conseil régional Provence Alpes-Côte d’Azur, Conseil Départemental 06, Cancéropôle PACA, Gis Ibisa and Inserm, the IRCAN’s Animal core facility, supported by FEDER, Région Provence Alpes-Côte d’Azur, Conseil Départemental 06 and Inserm, the IRCAN’s Histology core facility, supported by Région Provence Alpes-Côte d’Azur, Cancéropôle PACA, and Université Côte d’Azur and, the University’s CCMA Electron Microscopy facility supported by Université Côte d’Azur, Région Provence Alpes-Côte d’Azur, Conseil Départemental 06, and Gis Ibisa.

This work was made possible by grants to VP-F from the ANR (Agence Nationale de la Recherche) ANR-16-CE16-0024-01.

## Author contributions

ECG and SB designed, performed experiments and analyzed data on cellular and mouse models, and contributed to the main text. BR, AM-C, FL, GA and CC performed and analyzed experiments on mouse models and patient fibroblasts. KF designed, performed experiments and analyzed data on BN-PAGE experiments with SB. SL-G performed electron microscopy experiments. VP-F designed, supervised the project and drafted the manuscript. All of the authors read and approved the final submission.

---

**Supplementary Figure 1. Negative controls for PLA analysis.** Negative control experiments, using one antibody only, were performed in parallel to check the absence of PLA signal. P1, patient fibroblasts.

**Supplementary Figure 2. A.** Control (C) or patient (P1, P2) fibroblasts were transfected with scrambled siRNA (siScr), as negative control, or SLP2 siRNA (siSLP2) for experiments reported in Fig.4A-D. Representative western blot of SLP2 expression level 72h after transfection. Antibodies against HSP60 and GAPDH were used for loading control**. B.** Control (C) or patient (P1, P2) fibroblasts were transfected with scrambled siRNA (siScr), as negative control, or OMA1 siRNA (siOMA1) for experiments reported in Fig.4E-H. Representative western blot of OMA1 expression level 72h after transfection. Antibodies against HSP60 and GAPDH were used for loading control.

**Supplementary Figure 3. A.** Control (C) or patient (P1, P2) fibroblasts were transfected with scrambled siRNA (siScr), as negative control, or OMA1 siRNA (siOMA1) for experiments reported in Fig.5D-E. Representative western blot of OMA1 expression level 72h after transfection. Antibodies against HSP60 and GAPDH were used for loading control. **B.** Control (C) or patient (P1, P2) fibroblasts were transfected with scrambled siRNA (siScr), as negative control, or PHB2 siRNA (siPHB2) for experiments reported in Fig.5F. Representative western blot of PHB2 expression level 72h after transfection. Antibodies against HSP60 and GAPDH were used for loading control.

**Supplementary Figure 4. A.** Control (C) or patient (P1, P2) fibroblasts were transfected with scrambled siRNA (siScr), as negative control, or PINK1 siRNA (siPINK1) for experiments reported in Fig.6A-B. RT-qPCR analysis of *PINK1* in siRNA-transfected fibroblasts. Results were shown as fold change of *PINK1* mRNA expression in siPINK1 transfected fibroblasts relative to siScr transfected fibroblasts. Data were normalized to beta-2 macroglobulin (B2M) or 28S. Results shown are mean ± SEM from 2 independent experiments. **B.** Body weight curves showing failure to gain weight from 10 weeks of age for male and from 21 weeks for female KI (*Chchd10^S59L/+^*) *mito-*QC mice (arrows) that were significantly leaner than their littermates. Shown are mean±SD. **C.** Survival curve of the KI (*Chchd10^S59L/+^*) *mito-*QC mice **D.** Western blot on total extracts performed after microdissection of the hippocampus from *Chchd10^S59L/+^* mice (S59L/+) and control littermates (+/+), using antibodies against PHB2 and HSP60 (as loading control).

**Supplementary Table 1. Antibodies used in the study.**

## References

Abdulla S, Machts J, Kaufmann J, Patrick K, Kollewe K, Dengler R, Heinze H-J, Petri S, Vielhaber S & Nestor PJ (2014) Hippocampal degeneration in patients with amyotrophic lateral sclerosis. Neurobiol Aging 35: 2639–2645

Acin-Perez R, Lechuga-Vieco AV, Del Mar Muñoz M, Nieto-Arellano R, Torroja C, Sánchez-Cabo F, Jiménez C, González-Guerra A, Carrascoso I, Benincá C, et al (2018) Ablation of the stress protease OMA1 protects against heart failure in mice. Sci Transl Med 10

Ajroud-Driss S, Fecto F, Ajroud K, Lalani I, Calvo SE, Mootha VK, Deng H-X, Siddique N, Tahmoush AJ, Heiman-Patterson TD, et al (2015) Mutation in the novel nuclear-encoded mitochondrial protein CHCHD10 in a family with autosomal dominant mitochondrial myopathy. Neurogenetics 16: 1–9

Akepati VR, Müller E-C, Otto A, Strauss HM, Portwich M & Alexander C (2008) Characterization of OPA1 isoforms isolated from mouse tissues. J Neurochem 106: 372–383

Anand R, Wai T, Baker MJ, Kladt N, Schauss AC, Rugarli E & Langer T (2014) The i-AAA protease YME1L and OMA1 cleave OPA1 to balance mitochondrial fusion and fission. J Cell Biol 204: 919–929

Anderson CJ, Bredvik K, Burstein SR, Davis C, Meadows SM, Dash J, Case L, Milner TA, Kawamata H, Zuberi A, et al (2019) ALS/FTD mutant CHCHD10 mice reveal a tissue-specific toxic gain-of-function and mitochondrial stress response. Acta Neuropathol 138: 103–121

Auranen M, Ylikallio E, Shcherbii M, Paetau A, Kiuru-Enari S, Toppila JP & Tyynismaa H (2015) CHCHD10 variant p.(Gly66Val) causes axonal Charcot-Marie-Tooth disease. Neurol Genet 1: e1

Baek M, Choe Y-J, Bannwarth S, Kim J, Maitra S, Dorn GW, Taylor JP, Paquis-Flucklinger V & Kim NC (2021) TDP-43 and PINK1 mediate CHCHD10S59L mutation-induced defects in Drosophila and in vitro. Nat Commun 12: 1924

Baker MJ, Lampe PA, Stojanovski D, Korwitz A, Anand R, Tatsuta T & Langer T (2014) Stress-induced OMA1 activation and autocatalytic turnover regulate OPA1-dependent mitochondrial dynamics. EMBO J 33: 578–593

Bannwarth S, Ait-El-Mkadem S, Chaussenot A, Genin EC, Lacas-Gervais S, Fragaki K, Berg-Alonso L, Kageyama Y, Serre V, Moore DG, et al (2014) A mitochondrial origin for frontotemporal dementia and amyotrophic lateral sclerosis through CHCHD10 involvement. Brain 137: 2329–2345

Bannwarth S, Figueroa A, Fragaki K, Destroismaisons L, Lacas-Gervais S, Lespinasse F, Vandenbos F, Pradelli LA, Ricci J-E, Rötig A, et al (2012) The human MSH5 (MutSHomolog 5) protein localizes to mitochondria and protects the mitochondrial genome from oxidative damage. Mitochondrion 12: 654–665

Barrera M, Koob S, Dikov D, Vogel F & Reichert AS (2016) OPA1 functionally interacts with MIC60 but is dispensable for crista junction formation. FEBS Lett 590: 3309–3322

Bloemberg D & Quadrilatero J (2019) Autophagy, apoptosis, and mitochondria: molecular integration and physiological relevance in skeletal muscle. Am J Physiol Cell Physiol 317: C111–C130

Chen Y, Deng J, Wang P, Yang M, Chen X, Zhu L, Liu J, Lu B, Shen Y, Fushimi K, et al (2016) PINK1 and Parkin are genetic modifiers for FUS-induced neurodegeneration. Hum Mol Genet 25: 5059–5068

Christie DA, Lemke CD, Elias IM, Chau LA, Kirchhof MG, Li B, Ball EH, Dunn SD, Hatch GM & Madrenas J (2011) Stomatin-like protein 2 binds cardiolipin and regulates mitochondrial biogenesis and function. Mol Cell Biol 31: 3845–3856

Christie DA, Mitsopoulos P, Blagih J, Dunn SD, St-Pierre J, Jones RG, Hatch GM & Madrenas J (2012) Stomatin-like protein 2 deficiency in T cells is associated with altered mitochondrial respiration and defective CD4+ T cell responses. J Immunol 189: 4349–4360

Cipolat S, Martins de Brito O, Dal Zilio B & Scorrano L (2004) OPA1 requires mitofusin 1 to promote mitochondrial fusion. Proc Natl Acad Sci U S A 101: 15927–15932

Cogliati S, Frezza C, Soriano ME, Varanita T, Quintana-Cabrera R, Corrado M, Cipolat S, Costa V, Casarin A, Gomes LC, et al (2013) Mitochondrial cristae shape determines respiratory chain supercomplexes assembly and respiratory efficiency. Cell 155: 160–171

Cozzolino M, Rossi S, Mirra A & Carrì MT (2015) Mitochondrial dynamism and the pathogenesis of Amyotrophic Lateral Sclerosis. Front Cell Neurosci 9: 31

Da Cruz S, Parone PA, Gonzalo P, Bienvenut WV, Tondera D, Jourdain A, Quadroni M & Martinou J-C (2008) SLP-2 interacts with prohibitins in the mitochondrial inner membrane and contributes to their stability. Biochim Biophys Acta 1783: 904–911

Deshwal S, Fiedler KU & Langer T (2020) Mitochondrial Proteases: Multifaceted Regulators of Mitochondrial Plasticity. Annu Rev Biochem 89: 501–528

Eriksson PS, Perfilieva E, Björk-Eriksson T, Alborn AM, Nordborg C, Peterson DA & Gage FH (1998) Neurogenesis in the adult human hippocampus. Nat Med 4: 1313–1317

Frezza C, Cipolat S, Martins de Brito O, Micaroni M, Beznoussenko GV, Rudka T, Bartoli D, Polishuck RS, Danial NN, De Strooper B, et al (2006) OPA1 controls apoptotic cristae remodeling independently from mitochondrial fusion. Cell 126: 177–189

Friedman JR, Mourier A, Yamada J, McCaffery JM & Nunnari J (2015) MICOS coordinates with respiratory complexes and lipids to establish mitochondrial inner membrane architecture. Elife 4

Gami-Patel P, van Dijken I, Meeter LH, Melhem S, Morrema THJ, Scheper W, van Swieten JC, Rozemuller AJM, Dijkstra AA & Hoozemans JJM (2021) Unfolded protein response activation in C9orf72 frontotemporal dementia is associated with dipeptide pathology and granulovacuolar degeneration in granule cells. Brain Pathol 31: 163–173

Genin EC, Bannwarth S, Lespinasse F, Ortega-Vila B, Fragaki K, Itoh K, Villa E, Lacas-Gervais S, Jokela M, Auranen M, et al (2018) Loss of MICOS complex integrity and mitochondrial damage, but not TDP-43 mitochondrial localisation, are likely associated with severity of CHCHD10-related diseases. Neurobiology of Disease 119: 159–171

Genin EC, Madji Hounoum B, Bannwarth S, Fragaki K, Lacas-Gervais S, Mauri-Crouzet A, Lespinasse F, Neveu J, Ropert B, Augé G, et al (2019) Mitochondrial defect in muscle precedes neuromuscular junction degeneration and motor neuron death in CHCHD10S59L/+ mouse. Acta Neuropathol 138: 123–145

Genin EC, Plutino M, Bannwarth S, Villa E, Cisneros-Barroso E, Roy M, Ortega-Vila B, Fragaki K, Lespinasse F, Pinero-Martos E, et al (2016) CHCHD10 mutations promote loss of mitochondrial cristae junctions with impaired mitochondrial genome maintenance and inhibition of apoptosis. EMBO Mol Med 8: 58–72

Glytsou C, Calvo E, Cogliati S, Mehrotra A, Anastasia I, Rigoni G, Raimondi A, Shintani N, Loureiro M, Vazquez J, et al (2016) Optic Atrophy 1 Is Epistatic to the Core MICOS Component MIC60 in Mitochondrial Cristae Shape Control. Cell Rep 17: 3024–3034

Gómez-Pinedo U, Villar-Quiles RN, Galán L, Matías-Guiu JA, Benito-Martin MS, Guerrero-Sola A, Moreno-Ramos T & Matías-Guiu J (2016) Immununochemical Markers of the Amyloid Cascade in the Hippocampus in Motor Neuron Diseases. Front Neurol 7: 195

Head B, Griparic L, Amiri M, Gandre-Babbe S & van der Bliek AM (2009) Inducible proteolytic inactivation of OPA1 mediated by the OMA1 protease in mammalian cells. J Cell Biol 187: 959–966

Hernando-Rodríguez B & Artal-Sanz M (2018) Mitochondrial Quality Control Mechanisms and the PHB (Prohibitin) Complex. Cells 7

Huang X, Wu BP, Nguyen D, Liu Y-T, Marani M, Hench J, Bénit P, Kozjak-Pavlovic V, Rustin P, Frank S, et al (2018) CHCHD2 accumulates in distressed mitochondria and facilitates oligomerization of CHCHD10. Hum Mol Genet 27: 3881–3900

Ishihara N, Fujita Y, Oka T & Mihara K (2006) Regulation of mitochondrial morphology through proteolytic cleavage of OPA1. EMBO J 25: 2966–2977

Johnson JO, Glynn SM, Gibbs JR, Nalls MA, Sabatelli M, Restagno G, Drory VE, Chiò A, Rogaeva E & Traynor BJ (2014) Mutations in the CHCHD10 gene are a common cause of familial amyotrophic lateral sclerosis. Brain 137: e311

Korwitz A, Merkwirth C, Richter-Dennerlein R, Tröder SE, Sprenger H-G, Quirós PM, López-Otín C, Rugarli EI & Langer T (2016) Loss of OMA1 delays neurodegeneration by preventing stress-induced OPA1 processing in mitochondria. J Cell Biol 212: 157–166

Lachén-Montes M, González-Morales A, Zelaya MV, Pérez-Valderrama E, Ausín K, Ferrer I, Fernández-Irigoyen J & Santamaría E (2017) Olfactory bulb neuroproteomics reveals a chronological perturbation of survival routes and a disruption of prohibitin complex during Alzheimer’s disease progression. Sci Rep 7: 9115

Liu T, Woo J-AA, Bukhari MZ, LePochat P, Chacko A, Selenica M-LB, Yan Y, Kotsiviras P, Buosi SC, Zhao X, et al (2020) CHCHD10-regulated OPA1-mitofilin complex mediates TDP-43-induced mitochondrial phenotypes associated with frontotemporal dementia. FASEB J 34: 8493–8509

Matsuda N, Sato S, Shiba K, Okatsu K, Saisho K, Gautier CA, Sou Y-S, Saiki S, Kawajiri S, Sato F, et al (2010) PINK1 stabilized by mitochondrial depolarization recruits Parkin to damaged mitochondria and activates latent Parkin for mitophagy. J Cell Biol 189: 211–221

McBride H & Soubannier V (2010) Mitochondrial function: OMA1 and OPA1, the grandmasters of mitochondrial health. Curr Biol 20: R274–276

McWilliams TG, Prescott AR, Allen GFG, Tamjar J, Munson MJ, Thomson C, Muqit MMK & Ganley IG (2016) mito-QC illuminates mitophagy and mitochondrial architecture in vivo. J Cell Biol 214: 333–345

Meeusen S, DeVay R, Block J, Cassidy-Stone A, Wayson S, McCaffery JM & Nunnari J (2006) Mitochondrial inner-membrane fusion and crista maintenance requires the dynamin-related GTPase Mgm1. Cell 127: 383–395

Merkwirth C, Dargazanli S, Tatsuta T, Geimer S, Löwer B, Wunderlich FT, von Kleist-Retzow J-C, Waisman A, Westermann B & Langer T (2008) Prohibitins control cell proliferation and apoptosis by regulating OPA1-dependent cristae morphogenesis in mitochondria. Genes Dev 22: 476–488

Merkwirth C, Martinelli P, Korwitz A, Morbin M, Brönneke HS, Jordan SD, Rugarli EI & Langer T (2012) Loss of prohibitin membrane scaffolds impairs mitochondrial architecture and leads to tau hyperphosphorylation and neurodegeneration. PLoS Genet 8: e1003021

Mitsopoulos P, Chang Y-H, Wai T, König T, Dunn SD, Langer T & Madrenas J (2015) Stomatin-like protein 2 is required for in vivo mitochondrial respiratory chain supercomplex formation and optimal cell function. Mol Cell Biol 35: 1838–1847

Montava-Garriga L, Singh F, Ball G & Ganley IG (2020) Semi-automated quantitation of mitophagy in cells and tissues. Mech Ageing Dev 185: 111196

Müller K, Andersen PM, Hübers A, Marroquin N, Volk AE, Danzer KM, Meitinger T, Ludolph AC, Strom TM & Weishaupt JH (2014) Two novel mutations in conserved codons indicate that CHCHD10 is a gene associated with motor neuron disease. Brain 137: e309

Narendra DP, Jin SM, Tanaka A, Suen D-F, Gautier CA, Shen J, Cookson MR & Youle RJ (2010) PINK1 is selectively stabilized on impaired mitochondria to activate Parkin. PLoS Biol 8: e1000298

Olichon A, Baricault L, Gas N, Guillou E, Valette A, Belenguer P & Lenaers G (2003) Loss of OPA1 perturbates the mitochondrial inner membrane structure and integrity, leading to cytochrome c release and apoptosis. J Biol Chem 278: 7743–7746

Osman C, Merkwirth C & Langer T (2009) Prohibitins and the functional compartmentalization of mitochondrial membranes. J Cell Sci 122: 3823–3830

Palomo GM, Granatiero V, Kawamata H, Konrad C, Kim M, Arreguin AJ, Zhao D, Milner TA & Manfredi G (2018) Parkin is a disease modifier in the mutant SOD1 mouse model of ALS. EMBO Mol Med 10

Penttilä S, Jokela M, Bouquin H, Saukkonen AM, Toivanen J & Udd B (2015) Late onset spinal motor neuronopathy is caused by mutation in CHCHD10. Ann Neurol 77: 163–172

Pfanner N, Laan M van der, Amati P, Capaldi RA, Caudy AA, Chacinska A, Darshi M, Deckers M, Hoppins S, Icho T, et al (2014) Uniform nomenclature for the mitochondrial contact site and cristae organizing system. J Cell Biol 204: 1083–1086

Pickrell AM & Youle RJ (2015) The roles of PINK1, parkin, and mitochondrial fidelity in Parkinson’s disease. Neuron 85: 257–273

Quirós PM, Langer T & López-Otín C (2015) New roles for mitochondrial proteases in health, ageing and disease. Nat Rev Mol Cell Biol 16: 345–359

Raaphorst J, van Tol MJ, de Visser M, van der Kooi AJ, Majoie CB, van den Berg LH, Schmand B & Veltman DJ (2015) Prose memory impairment in amyotrophic lateral sclerosis patients is related to hippocampus volume. Eur J Neurol 22: 547–554

Salmon MichaelA, Esiri MargaretM & Ruderman NeilB (1971) Myopathic disorder associated with mitochondrial abnormalities, hyperglycaemia, and hyperketonaemia. The Lancet 298: 290–293

Signorile A, Sgaramella G, Bellomo F & De Rasmo D (2019) Prohibitins: A Critical Role in Mitochondrial Functions and Implication in Diseases. Cells 8

Sirkis DW, Geier EG, Bonham LW, Karch CM & Yokoyama JS (2019) Recent advances in the genetics of frontotemporal dementia. Curr Genet Med Rep 7: 41–52

Straub IR, Weraarpachai W & Shoubridge EA (2021) Multi-OMICS study of a CHCHD10 variant causing ALS demonstrates metabolic rewiring and activation of endoplasmic reticulum and mitochondrial unfolded protein responses. Hum Mol Genet

Sun X, Duan Y, Qin C, Li J-C, Duan G, Deng X, Ni J, Cao X, Xiang K, Tian K, et al (2018) Distinct multilevel misregulations of Parkin and PINK1 revealed in cell and animal models of TDP-43 proteinopathy. Cell Death Dis 9: 953

Tondera D, Grandemange S, Jourdain A, Karbowski M, Mattenberger Y, Herzig S, Da Cruz S, Clerc P, Raschke I, Merkwirth C, et al (2009) SLP-2 is required for stress-induced mitochondrial hyperfusion. EMBO J 28: 1589–1600

Tsai P-I, Lin C-H, Hsieh C-H, Papakyrikos AM, Kim MJ, Napolioni V, Schoor C, Couthouis J, Wu R-M, Wszolek ZK, et al (2018) PINK1 Phosphorylates MIC60/Mitofilin to Control Structural Plasticity of Mitochondrial Crista Junctions. Mol Cell 69: 744–756.e6

Viana MP, Levytskyy RM, Anand R, Reichert AS & Khalimonchuk O (2021) Protease OMA1 modulates mitochondrial bioenergetics and ultrastructure through dynamic association with MICOS complex. iScience 24: 102119

Wai T, García-Prieto J, Baker MJ, Merkwirth C, Benit P, Rustin P, Rupérez FJ, Barbas C, Ibañez B & Langer T (2015) Imbalanced OPA1 processing and mitochondrial fragmentation cause heart failure in mice. Science 350: aad0116

Wai T, Saita S, Nolte H, Müller S, König T, Richter-Dennerlein R, Sprenger H-G, Madrenas J, Mühlmeister M, Brandt U, et al (2016) The membrane scaffold SLP2 anchors a proteolytic hub in mitochondria containing PARL and the i-AAA protease YME1L. EMBO Rep 17: 1844–1856

Wang T, Liu H, Itoh K, Oh S, Zhao L, Murata D, Sesaki H, Hartung T, Na CH & Wang J (2021) C9orf72 regulates energy homeostasis by stabilizing mitochondrial complex I assembly. Cell Metab 33: 531–546.e9

Wei Y, Chiang W-C, Sumpter R, Mishra P & Levine B (2017) Prohibitin 2 Is an Inner Mitochondrial Membrane Mitophagy Receptor. Cell 168: 224–238.e10

Wu D, Jian C, Peng Q, Hou T, Wu K, Shang B, Zhao M, Wang Y, Zheng W, Ma Q, et al (2020) Prohibitin 2 deficiency impairs cardiac fatty acid oxidation and causes heart failure. Cell Death Dis 11: 181

Yan C, Gong L, Chen L, Xu M, Abou-Hamdan H, Tang M, Désaubry L & Song Z (2020) PHB2 (prohibitin 2) promotes PINK1-PRKN/Parkin-dependent mitophagy by the PARL-PGAM5-PINK1 axis. Autophagy 16: 419–434

